# ApoC-III overexpression and LDLr-/- protect mice from DSS-colitis: identifying a new role for lipoprotein metabolism in Tregs

**DOI:** 10.1101/823690

**Authors:** Cayla N Rodia, Diana Li, Nicholas S Tambini, Zania K Johnson, Evan R Jellison, Anthony T Vella, Alison B Kohan

## Abstract

**Objective:** Cellular metabolism is a key regulator of CD4^+^Foxp3^+^ regulatory T cell (Treg) homeostasis, but the foundational studies in this area use free fatty acid treatment as a proxy for plasma triglycerides. In vivo, plasma triglyceride is the main source of fatty acids for cells, not free fatty acids.

**Design/Results:** Using apolipoprotein C-III transgenic and LDLr^-/-^ mice, we report that the loss of lipoprotein triglycerides transport in these models results in protection from DSS-colitis and accumulation of intestinal Tregs and plasma IL-10. Total loss of apoC-III increases colitis severity. Tregs exposed to apoC-III increase lipolysis and fatty acid oxidation and apoC-III inhibits Bodipy-triglyceride uptake. Therapeutic treatment of WT mice with apoC-III-containing lipoproteins protects mice from colitis.

**Conclusion:** Our data suggest that therapies that reduce apoC-III could have negative effects in patients who are at risk of IBD, and conversely, that apoC-III could be a new therapeutic target to stimulate intestinal Tregs and IL-10 for the management of IBD. These data identify apoC-III and lipoprotein metabolism as a novel regulator of tolerance in the intestine.

**Summary Box:** * *What is already known about this subject*:

▪ The relative capacity to use either glucose or FFA to generate acetyl CoA for mitochondrial fatty acid oxidation is a critical driver of Treg and T cell activity and proliferation.
▪ ApoC-III is a known regulator of triglyceride and fatty acid metabolism in cells via LPL and LDLr endocytosis pathways
▪ ApoC-III is reduced in Crohn’s and Colitis patients.

* *What are the new findings*:

▪ We show that Tregs express triglyceride transporters, and that LDLr expression is enriched in Tregs from the mesenteric lymph nodes.
▪ We show that T cells are capable of endocytosing triglyceride from lipoproteins, and this process is inhibited by apoC-III.
▪ Tregs from *apoC-III^Tg^* are metabolically unique from *WT* Tregs and they upregulate the genes of lipolysis, and have an increase in basal respiration.
▪ The inhibition of TAG endocytosis, using 2 different models (LDLr^KO^ and apoC-III-transgenic mice), protects mice from colitis and stimulates the accumulation of Tregs and IL-10 in the gut.
▪ Intraperitoneal delivery of apoC-III on chylomicrons protects *WT* mice from DSS colitis.

*\* How might it impact on clinical practice in the foreseeable future?*

▪ Due to the protective role apoC-III plays in these mouse models of colitis, IBD risk should be carefully considered before prescribing patient anti-apoC-III lipid-lowering therapies.

## Introduction

*In vivo,* fatty acids are delivered to cells in the form of triacylglycerol (TAG) in chylomicrons and very-low density lipoproteins (VLDL). These lipoproteins are synthesized in the intestine and liver, respectively, and carry TAG from dietary sources or from liver stores. Cells are able to utilize the TAG for fuel through two processes: the hydrolysis of TAG to free fatty acids (FFA) via membrane-bound lipoprotein lipase (LPL) and the subsequent transport of those FFAs into the cell, or alternatively, endocytosis of lipoprotein TAG via the low-density lipoprotein receptor (LDLr), and the subsequent intracellular hydrolysis, oxidation, or storage of those FFAs. The way that a cell encounters a lipid fuel (either as a FFA bound to albumin or as a holoparticle lipoprotein) as significant impacts on the subsequent metabolic events [1,2]. Though FFA bound to albumin are present in blood, these only reach 10% the total amount of lipoprotein TAG available for cellular fuel [3].

In addition to carrying TAG, chylomicrons and VLDL also carry apolipoprotein C-III (apoC-III) and other apolipoproteins that regulate cellular metabolism. ApoC-III is a key regulator of plasma TAG metabolism, through its inhibition of both LPL and LDLr [4,5]. High levels of apoC-III on plasma lipoproteins delay the rate of TAG clearance into cells and stimulates hypertriglyceridemia. In humans and mice, plasma apoC-III independently correlates with increased incidence and severity of cardiovascular disease (CVD)[6]. *APOC3* is recognized as a valuable clinical target for reducing plasma TAG and CVD risk and antisense and antibody therapies against hepatic *APOC3* are currently in advanced trials [7,8].

ApoC-III also plays a critical role in intestinal metabolism. Overexpression of *APOC3* inhibits the rate of dietary TAG absorption, chylomicron secretion, and results in the secretion of small, dense chylomicrons[9]. Chylomicron apoC-III acts at the basolateral surface of enterocytes to deprives enterocytes of TAG fuel [10]. Therefore, apoC-III plays a major role in regulating TAG metabolism in multiple tissues.

Recently, Haberman et al. have identified that *APOC3* in the ileum of pediatric Crohn’s Disease patients is significantly reduced and that this is one of the largest changes in gene expression between healthy and Crohn’s Disease patients [11]. Other human studies find reductions in *APOC3* and the *APOA1/A4/C3* gene cluster in Crohn’s Disease patients [12,13]. Additionally, deletion of *HNF4a* (which regulates *APOC3* and the *APOA1/A4/C3* gene cluster) in intestinal epithelial cells causes spontaneous intestinal inflammation in mice [14]. This is the first evidence that *APOC3*, and the lipoproteins on which it resides, may play a causal or preventative role in inflammatory bowel disease (IBD).

In humans, IBD is exhibited by an imbalance between effector and tolerogenic T cells in the gut. CD4^+^CD25^+^Foxp3^+^ regulatory T cells (Tregs) are particularly critical to maintaining intestinal tolerance, through their suppression of inflammation via interleukin (IL)-10 secretion, whereas effector CD4^+^T cells secrete inflammatory tumor necrosis factor (TNF)-*α* [15,16]. In humans, the loss of Tregs results in severe refractory intestinal inflammation, and in mice the loss of Tregs or their IL-10 results in spontaneous colitis[17,18]. Most IBD patients become resistant to standard therapies, which primarily target the inflammatory arm of the disease, and one third of IBD patients do not achieve remission without bowel resection [19]. Thus, there is a major unmet clinical need to determine mechanisms for increasing the tolerogenic, Treg, and IL-10 driven arm of IBD.

Cellular metabolism is known regulator of Treg proliferation, differentiation, and cytokine secretion. Foundational studies have shown that metabolic fuel preference, and the capacity to use either glucose or FFAs for mitochondrial fatty acid oxidation (FAO), is a critical driver of immune cell function[20–22]. The limitation of these studies is that fatty acids are provided as palmitate bound to DMSO or bovine serum albumin (BSA), which does not approximate physiological TAG and fatty acid delivery via chylomicrons and VLDL. Whether Tregs acquire lipid substrates from circulating lipoproteins, and whether apoC-III, as a canonical regulator of this activity, influences Treg cellular metabolism and proliferation has never been addressed.

We demonstrate here that apoC-III transgenic (*apoC-III^Tg^*) mice accumulate intestinal Tregs and IL-10 and are protected from DSS-induced colitis. Overexpression of apoC-III shifts Treg metabolism towards lipolysis and FAO. We show that triglyceride-rich lipoproteins can deliver fatty acids to Jurkat T cells, and this is inhibited by apoC-III. We also show that *LDLr^-/-^* mice are protected from dextran sodium sulfate (DSS)-induced colitis and accumulate intestinal tolerogenic Tregs, and finally that IP injection of apoC-III-containing lipoproteins can protect *WT* mice from colitis. Our data suggest a new model: the inhibition of TAG delivery from lipoproteins to immune cells has profound effects on the accumulation of Tregs in the intestine and can protect mice from colitis in several different models. Our data suggest that therapies that reduce apoC-III, or LDLr, could have negative effects in patients who are at risk of IBD, and conversely, that apoC-III could improve IBD through a stimulation of intestinal Tregs and IL-10. These data identify apoC-III and lipoprotein metabolism as a novel regulator of tolerance in the intestine and as a potential new therapeutic target to stimulate intestinal Tregs and IL-10 for the management of IBD.

## Results

### Complete loss of APOC3 increases colitis severity and overexpression of APOC3 protects against colitis

Due to its role in raising plasma triglycerides and increasing CVD risk, apoC-III is a target for pharmaceutical inhibition. In IBD patients *APOC3* expression in the ileum is significantly reduced. We examined the effects of the total loss of *APOC3* in *apoC-III^-/-^* mice. We used the DSS model of colitis to determine whether *apoC-III^-/-^* mice would be protected from colitis. DSS-induced colitis is a well-characterized model of IBD and involves multiple immune responses including Tregs, effector T cells, TNF-α, and IFN-γ [23]. We exposed *WT* and *apoC-III^-/-^* mice to 4% DSS for 5 days followed by 5 days of water. We assessed the severity of colitis by qualitatively assessing disease progression with the macroscopic damage score. Body weight loss and colon lengths were comparable between *WT* and *apoC-III^-/-^* mice after DSS treatment (Figure 1, A and B). However, *apoC-III^-/-^* mice had significantly more-severe macroscopic damage with disease scores of 6.818 ± 0.5849 versus 4.364 ± 0.8008 in *WT* mice (*\*P=0.0224*), as well as significantly increased spleen weight (Figure 1, C and D). *ApoC-III^-/-^* mice and their *WT* counterparts had similar expression of *TNF-α* and *IL-6* in the colon (Figure 1E). Colonic lamina propria (LP) and mesenteric lymph node (mLN)-resident Treg populations were similar between *WT* and *apoC-III^-/-^* mice after DSS treatment (Figure 1, F and G). *ApoC-III^-/-^* mice had significantly greater CD14^+^ monocyte infiltration in the mLN compared to *WT* mice (*\*P=0.0432*) (Figure 1H). Monocyte infiltration into the gut-associated lymphoid tissue can aggravate colonic inflammation [24]. Therefore, the loss of apoC-III in mice is accompanied by an increase severity of DSS-colitis, which aligns with the findings in humans that *APOC3* expression is lowered in IBD.

**Figure 1:**
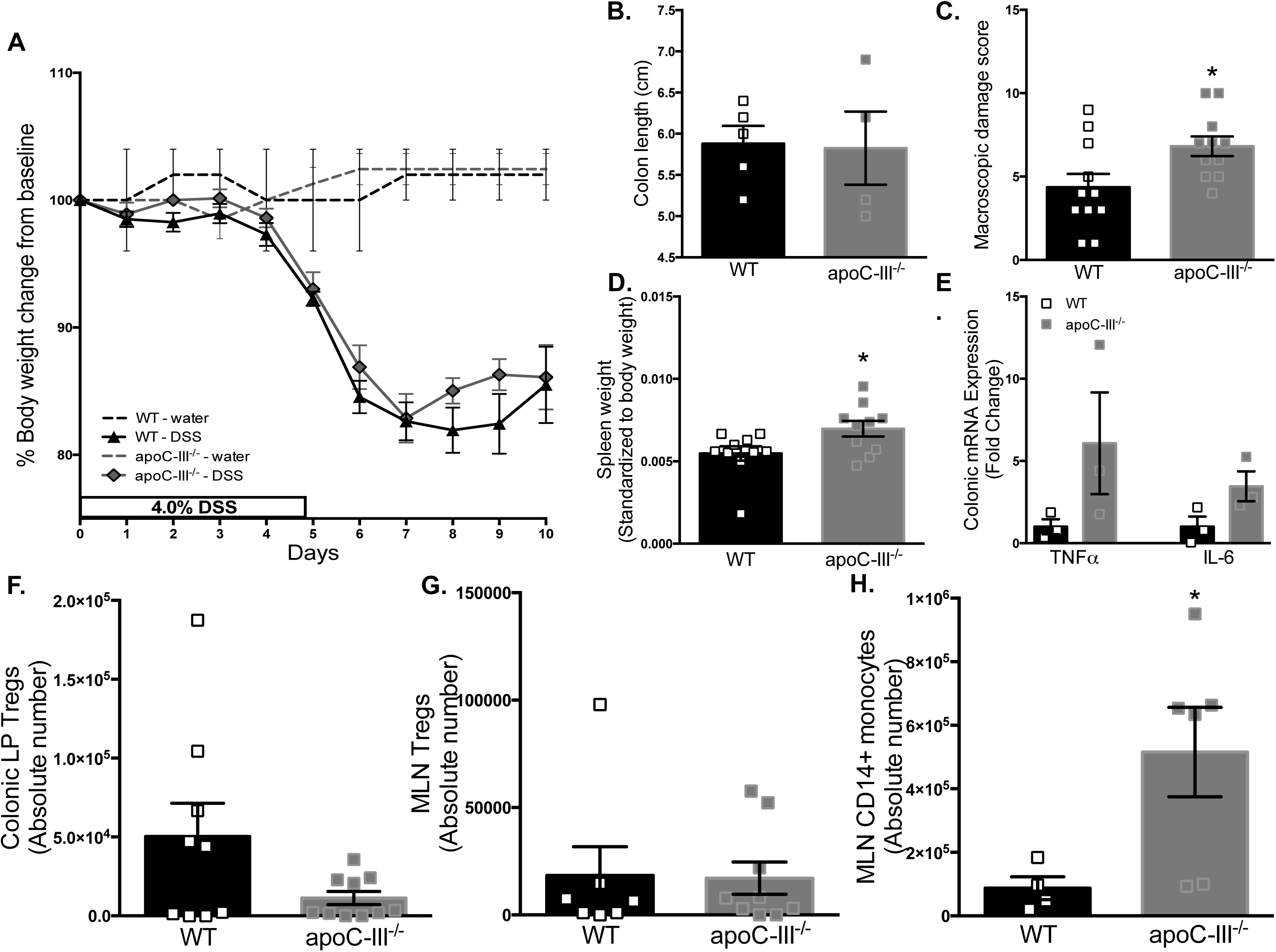
Loss of apoC-III increases colitis susceptibility. *ApoC-III^-/-^* and *WT* mice (aged 8 weeks), both maintained on the same C57Bl/6J background and on chow diet, were treated with 4% dextran sodium sulfate for 5 days followed by 5 days of water. **(A)** Percent body weight changes from baseline (before treatment). (n=9-12). Controls (given water only) are represented as dotted lines for each genotype. **B)** Colon length in centimeters (n=10) after treatment. **(C)** Macroscopic damage analysis of physiological symptoms of DSS. (n=11). The following data was collected after the complete treatment period (on Day 10): **(D)** Weight of spleen, standardized to mouse body weight. (n=4-5). **(E)** Colonic mRNA expression of *TNF-α* and *IL-6*. (n=3). After the 10 day treatment period, we used flow cytometric analysis to determine the absolute number of: **(F)** Colonic LP Tregs (n=9-10) **(G)** MLN Tregs. (n=6-10) and **(H)** MLN CD14^+^ monocytes. (n=5-6). Results are expressed as mean ± SEM, from three independent DSS treatments. For (A), 2way ANOVA (****P<0.0001 for genotype), with Sidak’s post-hoc analysis (no significant differences at specific days). Student’s T test, *P<0.05, comparison between genotypes.

We next asked whether overexpression of *APOC3* would protect mice from colitis. We used our colony of *apoC-III^Tg^* mice, which overexpress human *APOC3*, driven by the human *APOC3* promoter at ∼five-fold the level of endogenous mouse *APOC3* in liver, duodenum, and ileum. These mice maintain normal tissue-specific distribution and levels of mouse apoC-III (Supplementary Figure 1A)[25,26]. *ApoC-III^Tg^* mice also express the human transgene, as well as other apolipoprotein genes, in the colon (Supplementary Figure 1, A and B). *ApoC-III^Tg^* mice are hyperlipidemic: plasma triglycerides are ∼1,500 mg/dL and plasma cholesterol is ∼300 mg/dL (Supplementary Figure 1, C and D)*. ApoC-III^Tg^* mice have no obvious gut barrier disruption, as measured by plasma FITC-dextran after an oral gavage (Supplementary Figure 1E). We exposed *WT* and *apoC-III^Tg^* mice to 4% DSS for 5 days followed by 5 days of water. *ApoC-III^Tg^* mice lost significantly less weight than *WT* controls (11.5% compared to 15.7% weight loss, respectively; \*\*\*\**P<0.0001*) (Figure 2A). We found that *apoC-III^Tg^* mice were significantly protected from the physical symptoms of DSS including diarrhea, shivering, and lethargy compared to *WT* mice with a score of 0.7143 ± 0.1941 versus 4.571 ± 1.020, respectively (Figure 2B). *ApoC-III^Tg^* mice were resistant to colon shortening and thickening due to an accumulation of extracellular matrix, which is a well-established symptom of DSS-induced colitis (Figure 2C) [27]. *ApoC-III^Tg^* mice also showed significantly increased colon expression of the tight junction proteins occludin and zonulin-1 (*ZO-1*), strongly suggesting intact epithelial barrier function (Figure 1D) [28].

**Figure 2:**
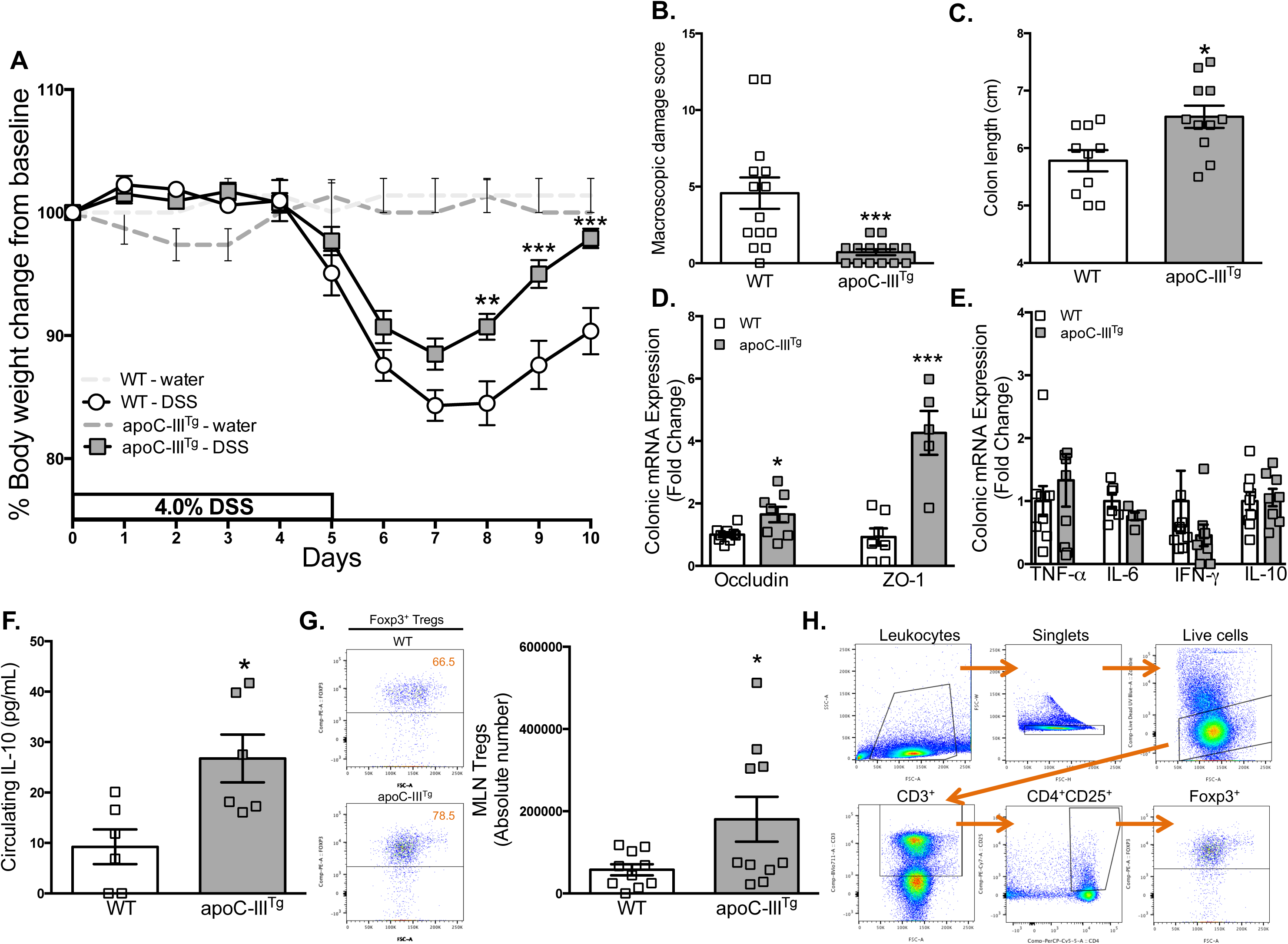
ApoC-III^Tg^ mice are protected from DSS-induced colitis. *ApoC-III^Tg^* and *WT* mice (aged 8 weeks), both maintained on the same C57Bl/6J background and on chow diet, were treated with 4% DSS for 5 days followed by 5 days of water. **(A)** Percent body weight changes from baseline (before treatment) (n=13-14). Controls (given water only) are represented as dotted lines for each genotype. **(B)** Macroscopic damage analysis of physiological symptoms of DSS. (n=14). The following data was collected after the complete treatment period (on Day 10): **(C)** Colon length in centimeters (n=14). **(D)** Colonic mRNA expression of tight junction complex genes, occludin and zonulin-1 (ZO-1) (n=5-9). **(E)** Colonic cytokine expression of TNF-α, IL-6, IFN-γ and IL-10. (n=5-9) **(F)** Circulating plasma IL-10 concentration measured by ELISA. (n=6) **(G)** Flow cytometric analysis of MLN Tregs (n=10). **(H)** Sample flow cytometry gating strategy to identify MLN Foxp3+ Tregs. Results are expressed as mean ± SEM. Results and dot plots are representative of at least 3 independent DSS experiments. For (A), 2way ANOVA (****P<0.0001 for genotype), with Sidak’s post-hoc analysis, **P<0.01, ***P<0.001, for comparisons between *apoC-III^Tg^* and *WT* at specific days. For (B-H), Student’s T test, *P<0.05, ***P<0.001, for comparisons between *apoC-III^Tg^* and *WT*.

After DSS treatment, *apoC-III^Tg^* and *WT* mice expressed comparable mRNA levels of *TNF-α, IL-6*, *IFN-γ,* and *IL-10* in the colon (Figure 1E). TNF-α concentrations at baseline (untreated, chow-fed) did not differ between *WT* and *apoC-III^Tg^* mice (0.9307 pg/mL versus 1.397 pg/mL, respectively; data not shown), and both genotypes experienced a non-significant change in plasma TNF-α after DSS treatment (Supplementary Figure 2A). An important inflammatory mediator of colitis is IL-17, which works in opposition to Treg IL-10 [29]. The mRNA expression level of this cytokine was not different at baseline (data not shown) or after the DSS treatment period between *WT* and *apoC-III^Tg^* mice (Supplementary Figure 2B). After DSS treatment, *apoC-III^Tg^* mice maintained significantly higher levels of circulating IL-10 (Figure 2F). Thus, *apoC-III^Tg^* mice showed resistance to DSS-induced colitis and maintained healthy intestinal barrier function.

The loss of IL-10-producing gut immune cells, such as CD103^+^ dendritic cells (DCs) and Tregs has been linked to symptoms of IBD[16]. We hypothesized that the resistance to DSS-induced colitis observed in *apoC-III^Tg^* mice was due to the accumulation of gut Tregs, which are potent sources of tolerogenic IL-10. We found that CD103 + DCs and mRNA expression of the Treg-related genes transforming growth factor (TGF)-β and FOXP3 are similar in the colon of both *WT* and *apoC-III^Tg^* mice (Supplementary Figure 2, C and D). Colonic LP Tregs in *apoC-III^Tg^* mice averaged 41,987 ± 22,137 cells versus 24,954 ± 15,949 in the colon of *WT* mice, although this difference was not significant (Supplementary Fig. 2, E and F). *ApoC-III^Tg^* mice also had similar colonic LP CD14^+^ monocytes compared to *WT* mice after DSS treatment (Supplementary Figure 2G). *ApoC-III^Tg^* and *WT* mice had comparable CD103^+^ DCs in the mLN (Supplementary Figure 2H), but *apoC-III^Tg^* mice had a significantly greater number of mLN-resident Tregs (*\*P=0.0124*; Figure 2, G and H). Thus, *APOC3* overexpression results in an accumulation of Tregs and IL-10 in the gut and protects mice from DSS-induced colitis.

### ApoC-III^Tg^ mice accumulate intestine-homing Tregs

Tregs in the gut, specifically the small intestinal and colonic LP and the gut-draining mLN, are potent sources of IL-10 [30]. We hypothesized that Treg accumulation in the gut of *apoC-III^Tg^* mice would also change the circulating cytokine profile. Plasma cytokine analysis between *WT* and *apoC-III^Tg^* mice showed no significant differences in circulating levels of tumor necrosis factor alpha TNF-α, IL-6, IL-4, or interferon gamma (IFN-γ) (Figure 3A). However, *apoC-III^Tg^* mice had significantly higher concentrations of anti-inflammatory circulating IL-10 compared to *WT* mice (*\*\*\*\*P<0.0001*; Figure 3B). We found no significant differences in intracellular IL-10 in Tregs from the spleen or mLN in both genotypes (Figure 3C).

**Figure 3:**
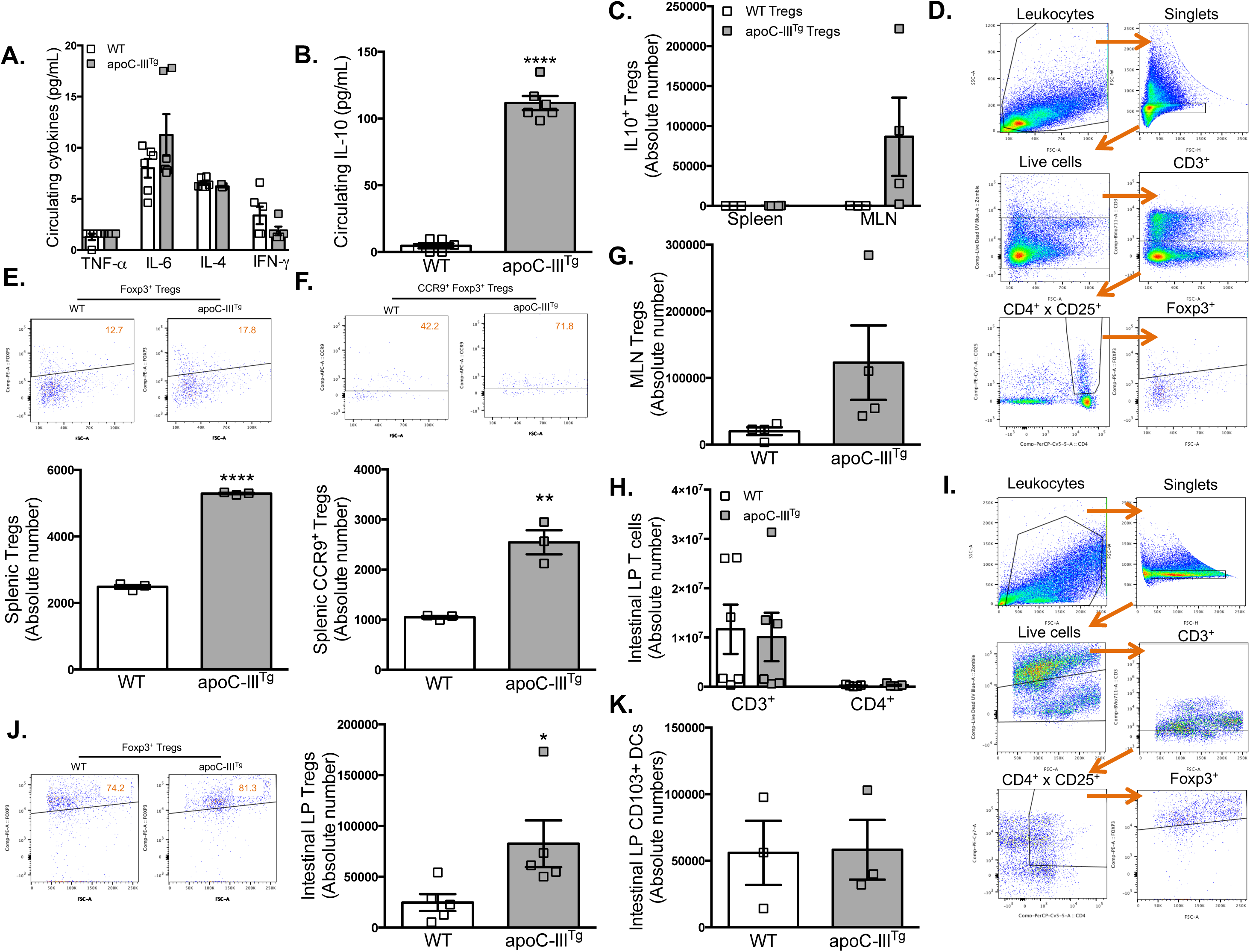
ApoC-III^Tg^ mice accumulate intestine-homing Tregs. *ApoC-III^Tg^* and *WT* mice, both maintained on the same C57Bl/6J background, were co-housed and maintained on chow diet. **(A)** Plasma cytokine concentrations measured by Multiplex analysis (n=4-6). **(B)** Basal plasma IL-10 concentration measured by ELISA. **(C)** Identification of IL-10^+^ Tregs in spleen and MLN measured by flow cytometry. (n=3-4)**. (D)** Sample gating strategy to identify splenic Foxp3+ Tregs by flow cytometry. We used flow cytometric analysis to identify the following: **(E)** Absolute number of CD4^+^CD25^+^Foxp3^+^ Tregs in spleen with representative dot plots for each genotype. (n=3-4) **(F)** Absolute number of splenic CCR9^+^ Tregs with representative dot plots for each genotype. (n=3-4) **(G)** Absolute number of MLN-resident CD4^+^CD25^+^Foxp3^+^ Tregs. (n=4) **(H)** Absolute number of Intestinal LP parent CD3+ T cells and CD4+ Helper T cells. (n=5-6) **(I)** Sample flow cytometry gating strategy to identify intestinal LP-Tregs. **(J)** Flow cytometric analysis of the absolute number of Intestinal LP-resident CD4^+^CD25^+^Foxp3^+^ Tregs with representative dot plots for each genotype. (n=5) **(K)** Absolute number of CD103^+^ DCs in intestinal LP (n=3). Dot plots are representative of at least three individual experiments. Results are expressed as mean ± SEM. Student’s T test, *P<0.05, **P<0.01, ****P<0.0001.

We found that *apoC-III^Tg^* mice also have an accumulation of splenic Tregs, compared to *WT* mice (*\*\*\*\*P<0.0001*; Figure 3, D and E). Tregs in *apoC-III^Tg^* spleen express C-C chemokine receptor type 9 (CCR9), which is activated by intestinal CCL25, an intestinal homing molecule (Figure 3F) [31]. This result suggests that splenic Tregs in *apoC-III^Tg^* mice might be homing to the intestine. We found no significant differences in mLN Treg populations or intestinal LP parent T cells (CD3^+^ or CD4^+^ Helper T cells) (Figure 3, G and H). In the intestinal LP, *apoC-III^Tg^* mice had a significant accumulation of Tregs compared to *WT* counterparts (Figure 3, I and J). This accumulation also occurred in the absence of changes to CD103^+^ DCs (Figure 3K).

We comprehensively analyzed the tissue compartments where apoC-III action is most potent (circulation and liver) to determine if excess apoC-III caused any other immune perturbations. We found no differences between *WT* and *apoC-III^Tg^* mice in the number of spleen-resident and circulating CD3^+^ T cells, CD4^+^ T cells, CD8α^+^ T cells, or CD19^+^ B cells (Supplementary Figure 3, A-D). We also observed no difference in splenic or circulating CD14^+^ monocytes (Supplementary Figure 3, E-H), despite previous reports showing that apoC-III-rich VLDLs can activate monocytes *in vitro* [32]. The liver, where apoC-III inhibits lipoprotein uptake, also had no differences in CD3^+^ T cells, CD4^+^ T cells, CD8α^+^ T cells, and CD25^+^ T cells (Supplementary Figure 4).

Because *apoC-III^Tg^* mice are hyperlipidemic (due to apoC-III inhibition of lipoprotein clearance from plasma to tissues) (Supplementary Figure 1C), we used another model of hyperlipidemia to evaluate whether raising plasma lipids would recapitulate the intestinal Treg phenotype. We fed *WT* mice a 12-week Western diet. This high-fat, high-cholesterol diet resulted in increased total plasma cholesterol and triglycerides in *WT* mice versus chow-fed controls, mimicking basal *apoC-III^Tg^* plasma lipids (Supplementary Figure 5, A and B). Flow cytometric analysis of the intestinal LP of Western diet-fed *WT* mice showed significant decreases in Treg populations compared to chow-fed controls (Supplementary Figure 5C). This finding suggests that hyperlipidemia alone is not the cause of increased Treg populations in *apoC-III^Tg^* mice.

### ApoC-III^Tg^ Tregs are phenotypically comparable to WT Tregs

We conducted an *ex vivo* analysis of *WT* and *apoC-III^Tg^* Tregs with and without T-cell receptor (TCR) stimulation to determine if there were phenotypic differences between these cells (Figure 4). Under both conditions, we found no differences in the levels of the Treg cell surface markers CTLA-4, PD-1, GITR, LAG3, and NP-1 between the genotypes (Figure 4A). *ApoC-III^Tg^* Tregs express the *APOC3* human transgene, but at a level 100 times less than liver expression of human *APOC3* (Supplementary Figure 1F). *WT* and *apoC-III^Tg^* Tregs have comparable expression levels of mouse *APOC3* (Supplementary Figure 1F). We stained Tregs with the cell cycle marker Ki-67 to identify which cells were actively proliferating[33]. Ki-67 staining was similar between *WT* and *apoC-III^Tg^* Tregs under both TCR-stimulated and unstimulated conditions (Figure 4B). To test whether isolated Tregs from *WT* and *apoC-III^Tg^* mice are different in their anergic state, we analyzed the amount of IL-2 in the cell culture media of isolated Tregs after 24 hours. Tregs are inherently anergic *in vitro* and require exogenous IL-2 to proliferate [34]. We found that both *WT* and *apoC-III^Tg^* Tregs are anergic and do not secrete any IL-2 (Figure 4C, shown as ND). From this, we can conclude that there are no major phenotypic differences between *WT* and *apoC-III^Tg^* Tregs excluding expression of the human *APOC3* transgene. This suggests that the change in Treg numbers in the intestine of *apoC-III^Tg^* mice is not due to a change in Treg activity or phenotype but is instead due to the accumulation of large numbers of Tregs in the gut.

**Figure 4:**
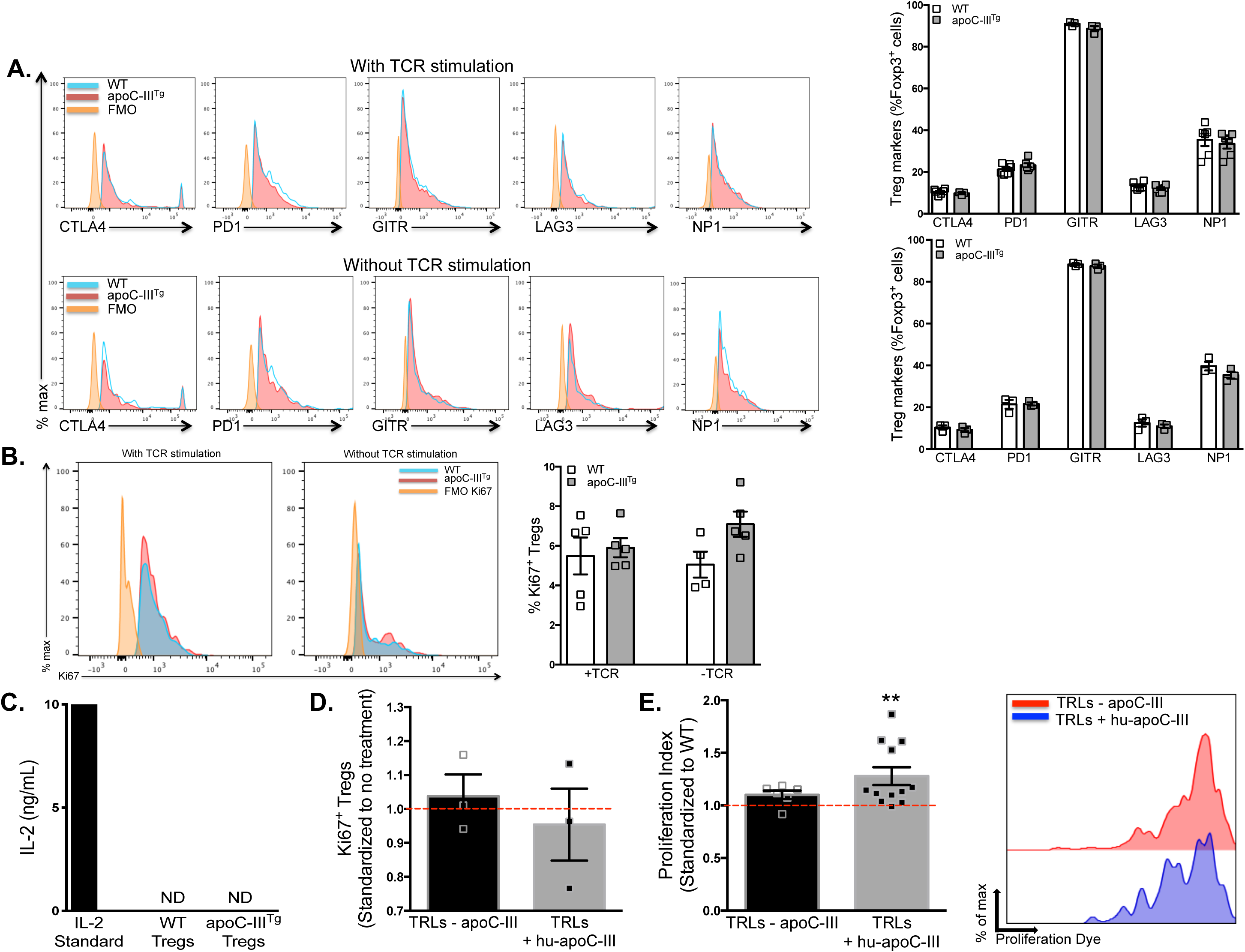
Tregs from apoC-III^Tg^ mice are phenotypically comparable to WT Tregs. *ApoC-III^Tg^ and WT* CD4^+^CD25^+^ Tregs were isolated and cultured 24 hours in complete media analyzed for: **(A)** Cell surface markers with TCR stimulation via anti-CD3 plus anti-CD28 for 24 hours (top row) and without TCR stimulation (bottom row) (n=3-6) and **(B)** proliferation marker Ki-67 with TCR stimulation (left) and without TCR stimulation (right) (n=3-6). **(C)** Cell culture media analysis of IL-2 concentration following 24 hour culture. **(D)** Ki-67 staining of *WT* CD4^+^CD25^+^ Tregs treated for 96 hours with TCR stimulation and with lipoproteins without apoC-III (TRLs – apoC-III) or lipoproteins with apoC-III (TRLs + hu-apoC-III), standardized to Tregs without treatment (represented as red line). **(E)** Jurkat T cell proliferation index measured by dilution of Tag It Violet proliferation dye in response to TRLs - apoC-III and TRLs + hu-apoC-III. Histogram is representative of Tag It Violet proliferation dye dilutions from four repeated experiments. (n=5 technical replicates per experiment). For **E**, -apoC-III and +hu-apoC-III treatments are standardized to treatment with lipoproteins with endogenous mouse levels of apoC-III (red dashed line). Results are expressed as mean ± SEM. Student’s T test, **P<0.01, comparison between TRLs - apoC-III and TRLs + hu-apoC-III treatments.

A possible explanation for the increase in intestinal Tregs in *apoC-III^Tg^* mice is that the parent CD4^+^ T cells are proliferating in response to apoC-III inhibition of lipid uptake. We used Jurkat T cells to model parent CD4^+^ T cells. A significant advantage of Jurkat T cells is that we can avoid sorting T cells via fluorescence-activated cell sorting (FACS), which can change the intracellular metabolism of the cells [35]. We stained Jurkat T cells with Tag It Violet™ proliferation dye and treated them with human apoC-III-containing triglyceride-rich lipoproteins (TRLs) or *apoC-III^-/-^* TRLs. Jurkat T cells treated with hu-apoC-III-containing TRLs had a greater proliferation index compared to cells treated with *apoC-III^-/-^* TRLs (TRLs-apoC-III) (Figure 4E). These data suggest that apoC-III, by inhibiting lipid uptake from lipoproteins to T cells, results in the proliferation of parent T cells, which might account for the increasing size of the intestinal Treg pool. This demonstrates that the modulation of substrate delivery from lipoproteins to T cells could be a critical regulator of their proliferation and metabolism.

### Tregs express lipid transporters, and apoC-III can inhibit their activity

For an immune cell to use circulating TAG, the lipid must go through key physiological events, including endocytosis by LDLr and hydrolysis of the fatty acids. Whether T cells and Tregs acquire TAG from circulating lipoproteins, and whether this influences their activity is unknown. We first determined that Tregs express lipid transporters by mRNA analysis from isolated splenic Tregs (purity >80%) from *WT* and *apoC-III^Tg^* mice (Figure 5A, Supplementary Figure 6, A and B). To assess if there are tissue-specific expression differences in LDLr on Tregs, we analyzed multiple anatomical locations including the spleen, the gut-draining mLN and intestinal LP of *WT* mice (Figure 5B). As a percentage of the total Treg population, 98.5% of intestinal LP Tregs express LDLr. This is in contrast to splenic Tregs, where only 72.58% of Tregs express this receptor. When comparing between genotypes, we found that *apoC-III^Tg^* Tregs from the intestinal LP express the highest levels of LDLr (Figure 5, C and D). This suggests that LDLr may be a functional marker of gut Tregs.

**Figure 5:**
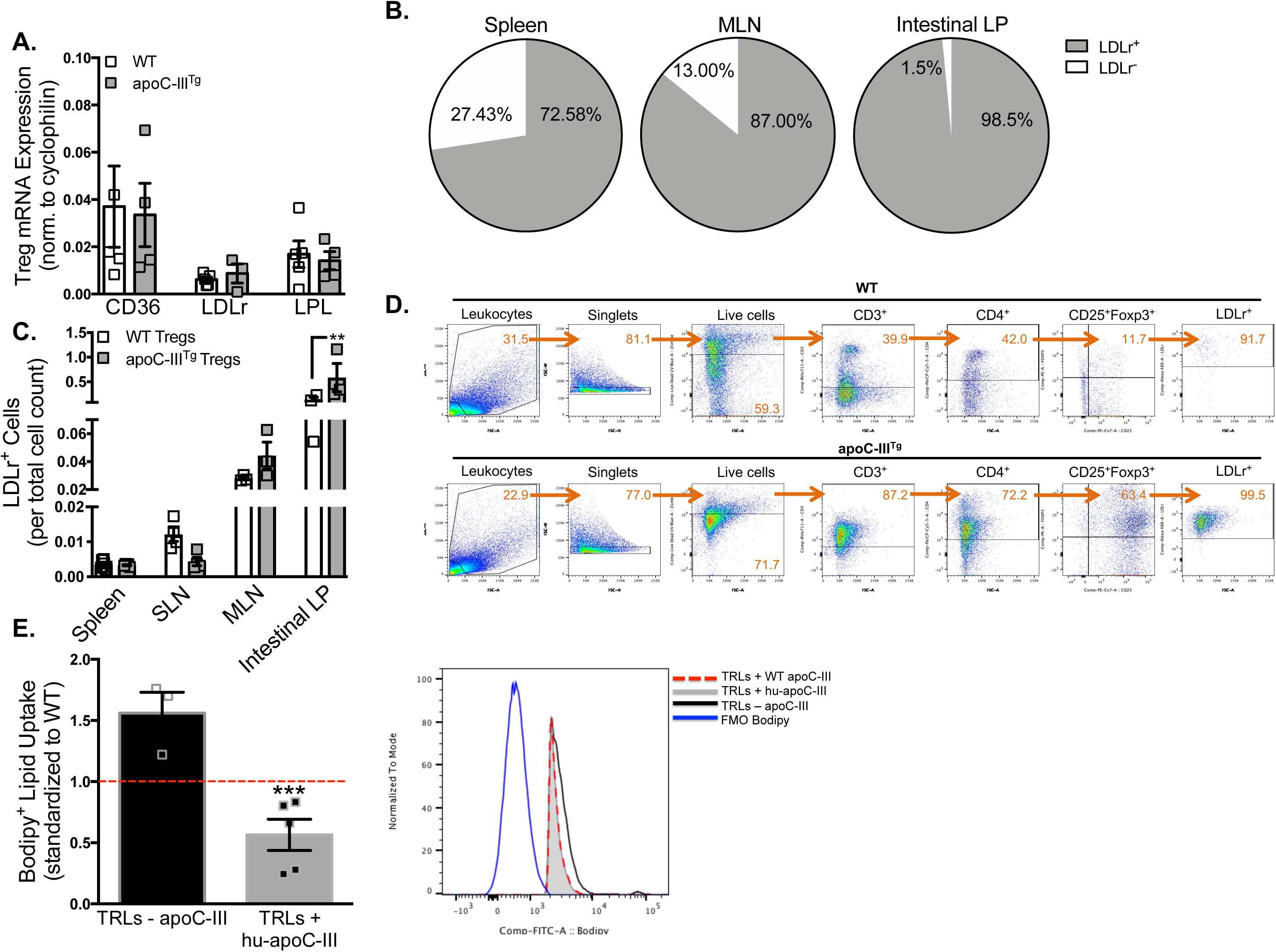
Tregs express lipid transporters, and apoC-III can inhibit their activity. *ApoC-III^Tg^* and *WT* mice, both maintained on the same C57Bl/6J background and on chow diet, were used for analysis of Tregs. **(A)** mRNA expression of lipid transporters: CD36, LDLr, and LPL in isolated CD4^+^CD25^+^ Tregs from *apoC-III^Tg^* and *WT* spleen. (n=3-5) **(B)** Ratio of LDLr^+^ and LDLr^-^ Tregs in spleen, MLN, and intestinal LP of *apoC-III^Tg^* and *WT* mice measured by flow cytometry. **(C)** Flow cytometric analysis of the absolute number of LDLr^+^ Tregs from the spleen, skin-draining lymph node (SLN), MLN, and intestinal LP of *apoC-III^Tg^* and *WT* mice, represented on a per Treg basis. (n=3-7) **(D)** Flow cytometry gating strategy to identify LDLr^+^ Tregs in the intestinal LP of *WT* mice (top row) and *apoC-III^Tg^* mice (bottom row). **(E)** Immortalized Jurkat T cells were cultured in complete RPMI-1640 media supplemented with 10% fetal bovine serum. Flow cytometric analysis was conducted on Jurkat T cells after various treatments to measured Bodipy-lipid fluorescence within cell. Treatment of Jurkat T cells with Bodipy-labeled lipoproteins containing no apoC-III (TRLs - apoC-III) and lipoproteins containing apoC-III (TRLs + hu-apoC-III). Histogram is representative Bodipy uptake from at least three repeated experiments. (n=3 technical replicates per experiment). Results are expressed as mean ± SEM. For **(C),** One-way ANOVA, with Sidak’s post-hoc analysis, **P<0.01, for comparisons between *apoC-III^Tg^* intestinal LP Tregs versus *WT* Intestinal LP Tregs. For **(E)**, treatments are standardized to treatment with lipoproteins with endogenous mouse levels of apoC-III (red dashed line). Student’s T test, ***P<0.001, comparison between TRLs - apoC-III and TRLs + hu-apoC-III treatments.

As proof of principle, we first determined whether naïve T cells can endocytose TAG from lipoproteins and whether this is inhibited by apoC-III. Previous studies have used fatty acids bound to BSA or DMSO, or have addressed the impact of oxidized lipids from LDL and HDL on immune cells [36–38]. First, we confirmed the expression of *LDLr* in Jurkat T cells, our model for parent T cells, after treating them with TRLs containing varying amounts of apoC-III (Supplementary Figure 7). *LDLr* expression in Jurkat T cells was similar between groups: no treatment (basal), treatment with TRLs from *WT* mice containing normal apoC-III levels, and treatment with TRLs containing excess apoC-III. Importantly, when treated with TRLs without apoC-III (no apoC-III) *LDLr* expression was downregulated which is a known regulatory step in liver in response to excess lipids [39].

To determine whether apoC-III acts extrinsically on Jurkat T cells to inhibit lipid uptake, we treated cells with Bodipy-labeled TRLs with varying amounts of apoC-III. When treated with Bodipy-labeled TRLs without apoC-III (TRLs-apoC-III), Jurkat T cells had the greatest lipid uptake compared to treatments with apoC-III (Figure 5E, Supplementary Figure 8A and B). The addition of human apoC-III to Bodipy-labeled TRLs (TRLs+hu-apoC-III) inhibited lipid uptake compared to those with no apoC-III, resulting in significantly decreased lipid uptake into the cells (Figure 5E). Thus, apoC-III inhibits lipid uptake from TRLs into Jurkat T cells in a cell-extrinsic manner just as apoC-III inhibits TAG uptake in other tissues that express LDLr [4].

### *In vivo* inhibition of triglyceride transport shifts the metabolic program of *apoC-III^Tg^* Tregs

Cellular metabolism is a key regulator of T cells and Tregs activity, including their proliferation at sites of inflammation, their differentiation to effector subsets, and their cytokine profile. We asked whether the inhibition TAG transport via LDLr had consequences for Treg metabolism, homeostasis, and activity *in vivo*. *ApoC-III^Tg^* Tregs have significantly increased expression of key lipolytic genes including hormone sensitive lipase (*HSL*) and patatin-like phospholipase domain (*PNPL*)-A2 (*\*\*\*P=0.0008*), and key FAO genes including carnitine palmitoyltransferase (*CPT*)1-β and acyl-coenzyme A oxidase (*ACOX*)1(\*\**P=0.0062*; Figure 6A). Concomitant with this increase in expression of lipolytic/oxidation genes, *apoC-III^Tg^* Tregs also showed decreased glucose transporter (*GLUT1)* mRNA compared to *WT* Tregs (\*\**P=0.0070*) (Figure 6A). This finding suggests a shift in *apoC-III^Tg^* Treg cellular metabolism from glycolysis and lipogenesis toward lipolysis and oxidative phosphorylation (OXPHOS). In contrast to *apoC-III^Tg^* Tregs, *apoC-III^-/-^* Tregs have decreased expression of *HSL*, which may suggest an alternative intracellular capacity for lipolysis (Figure 6B).

**Figure 6:**
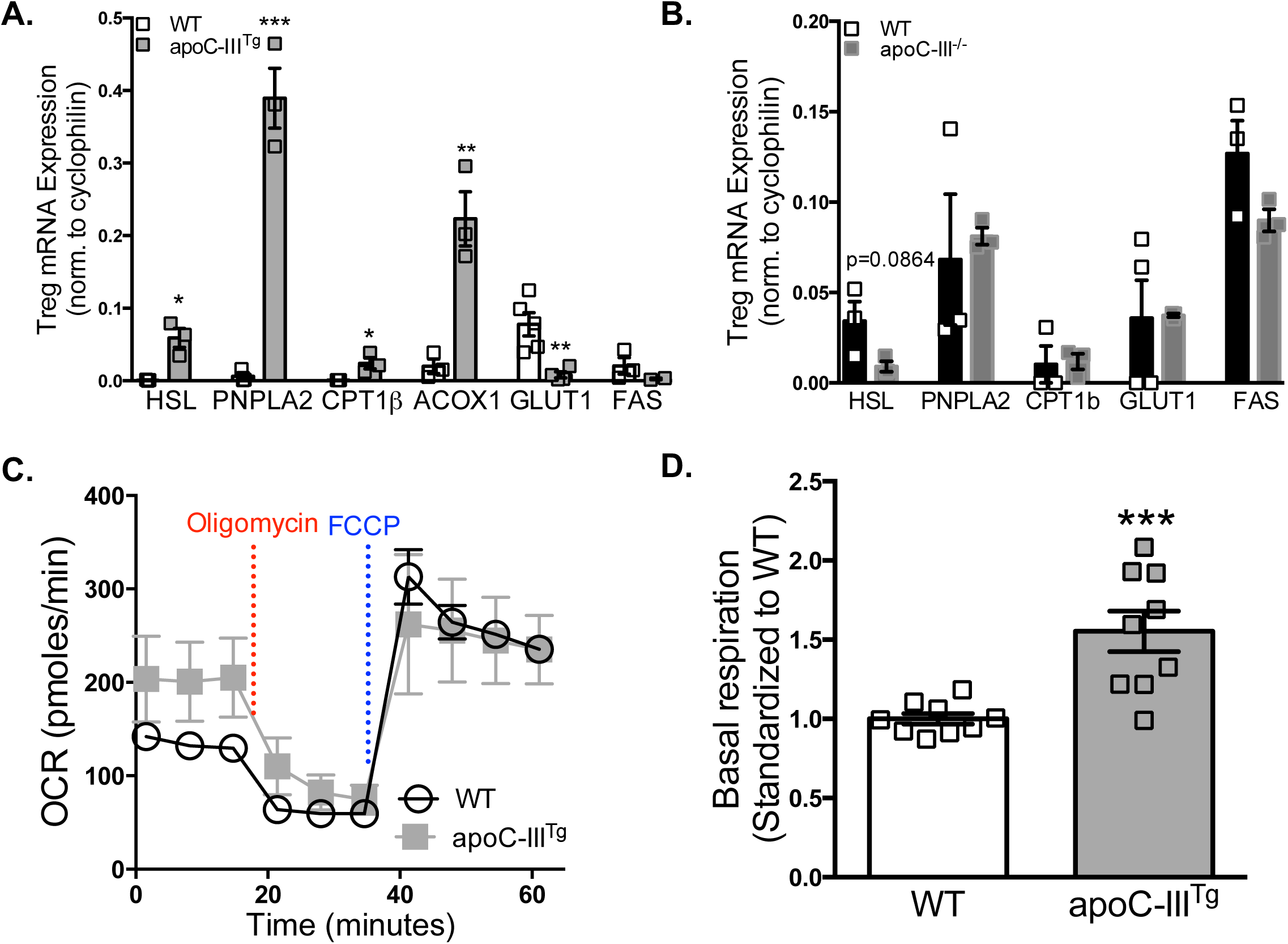
*In vivo*, inhibition of triglyeride transport shifts the metabolic program of apoC-III^Tg^ Tregs. Isolated CD4^+^CD25^+^ Treg from the spleen were used for the following: **(A)** Analysis of mRNA expression of metabolic genes comparing *WT* versus *apoC-III^Tg^* (n=3-4). **(B)** mRNA expression of metabolic genes comparing *WT* versus *apoC-III^-/-^* (n=3-4). **(C)** Seahorse metabolic flux analysis of oxygen consumption rate (OCR) of *apoC-III^Tg^* Tregs versus *WT* Tregs under basal conditions and in response to sequential treatment with Oligomycin and FCCP, and **(D)** Basal respiration standardized to *WT* Tregs. Three biological replicates were used in OCR analysis. Results are expressed as mean ± SEM. Student’s T test, *P<0.05, **P<0.01, ***P<0.001, for comparison between genotypes.

To confirm that this enzymatic capacity corresponded with a chance to *in vivo* metabolic flux, we used Seahorse analysis in splenic Tregs. We determined the oxygen consumption rate (OCR), which is a measure of OXPHOS, in isolated splenic Tregs from *WT* and *apoC-III^Tg^* mice (>80% purity) (Figure 6C). Although Tregs from *apoC-III^Tg^* and *WT* mice had similar responses to the ATP synthase inhibitor oligomycin and the proton gradient uncoupler FCCP, *apoC-III^Tg^* Tregs had an increased basal rate of OCR, confirming our gene expression analysis (Figure 6D). These data suggest that *apoC-III^Tg^* Tregs have increased basal respiration in response to excess apoC-III, and perhaps to the inhibited ability to take up lipid through LDLr. Although we show that Tregs upregulate lipolysis genes, it is unclear whether *apoC-III^Tg^* Tregs specifically break down intracellular TAG to fuel their increased basal respiration which is consistent with the mobilization of intracellular lipids in the absence of extracellular TAG transport.

### LDLr^-/-^ mice are protected against DSS-induced colitis

Since apoC-III’s main action is to inhibit LDLr, we next asked whether *LDLr^-/-^* mice would mirror colitis protection and recapitulate the tolerogenic phenotype seen in *apoC-III^Tg^* mice. To test this hypothesis, we treated *LDLr^-/-^* and *WT* mice with DSS to induce colitis (Figure 7). As predicted, the *LDLr^-/-^* mice lost less body weight over the course of the treatment period (*\*\*\*\*P<0.0001*; Figure 7A). Although colon lengths were comparable, the *LDLr^-/-^* mice had significantly lower macroscopic damage throughout the treatment period (Figure 7, B and C). *LDLr^-/-^* mice showed significant increases in CD4^+^ T cells and Tregs themselves in the mLN after DSS treatment (Figure 7, D and E, Supplementary Figure 9A). Although *LDLr^-/-^* mice had increased numbers of CD14^+^ monocytes infiltrating the colonic LP, they had significantly greater amounts of Tregs (Figure 7, F and G, Supplementary Figure 9B). This suggests that gut tolerance occurs in *LDLr^-/-^* mice as it does in *apoC-III^Tg^* mice during DSS treatment. Therefore, *LDLr^-/-^* mice phenocopied the *apoC-III^Tg^* mice colitis protection after DSS treatment. This supports our hypothesis that lipid uptake via LDLr, and its inhibition when lipoproteins contain apoC-III, results in protection from DSS-induced colitis, and corresponds with an accumulation of gut and MLN Tregs. This is a previously unexplored pathway to stimulate intestinal immune tolerance.

**Figure 7:**
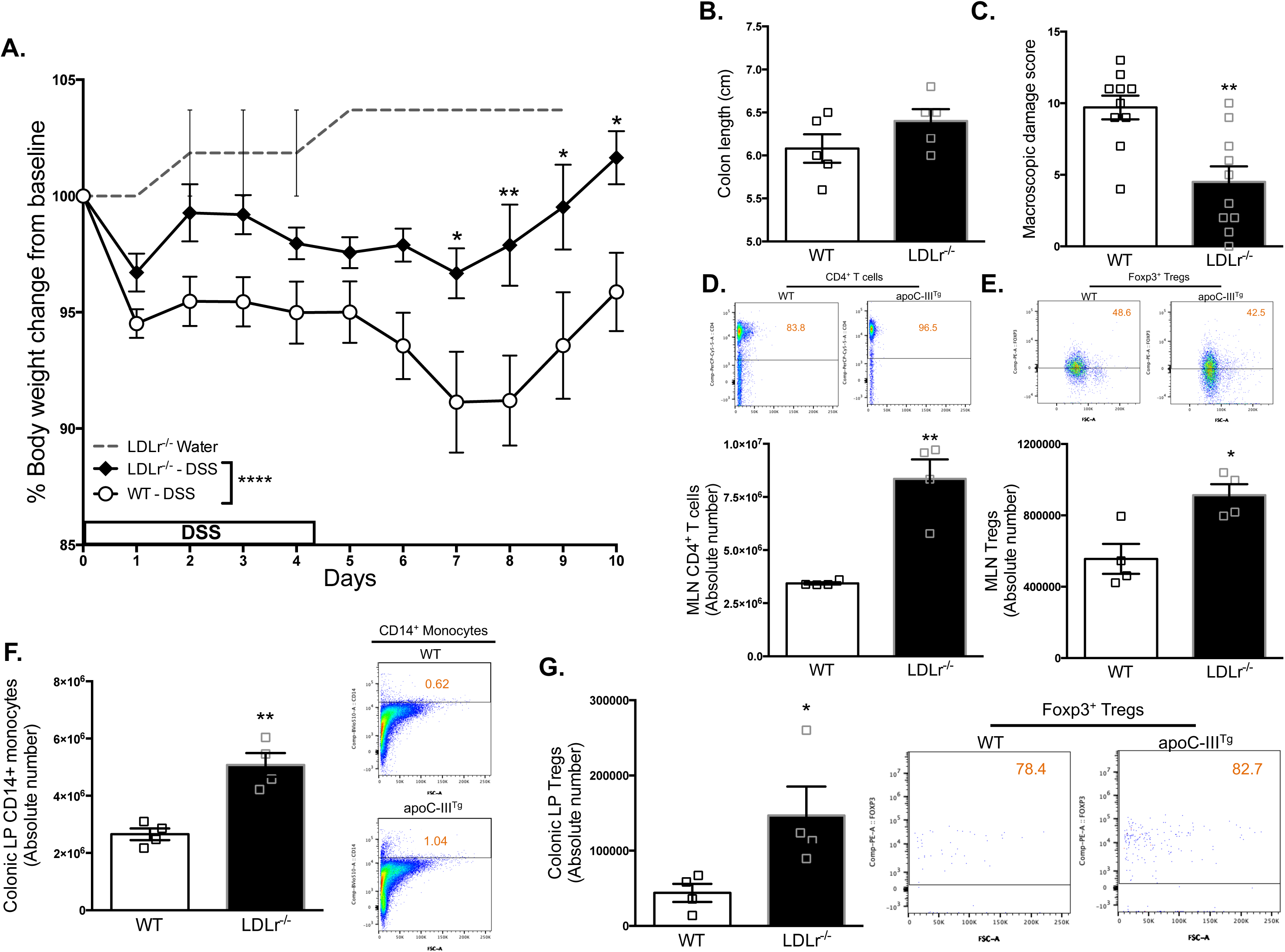
LDLr^-/-^ mice are protected against DSS-induced colitis. *LDLr^-/-^* and *WT* mice (aged 8 weeks), both maintained on the same C57Bl/6J background and on chow diet, were treated with DSS for 5 days followed by 5 days of water. **(A)** Percent body weight changes from baseline (before treatment). Controls (given water only) are represented as a dotted line for *LDLr^--/-^* only. (n=10) The following data was collected after the complete treatment period (on Day 10): **(B)** Colon length in centimeters (n=5). **(C)** Macroscopic damage analysis of physiological symptoms of DSS. (n=10). We used flow cytometric analysis to determine the absolute number of: **(D)** MLN CD4^+^ T cells, (n=5) **(E)** MLN Tregs, (n=5), **(F)** Colonic LP CD14^+^ monocytes, (n=5) and **(G)** Colonic LP Tregs. (n=5). All flow cytometric analysis is shown with representative dot plots for each genotype. Results are expressed as mean ± SEM, from at least 3 independent DSS experiments. For (A), 2way ANOVA (****P<0.0001 for genotype), with Sidak’s post-hoc analysis, *P<0.05, **P<0.01, **P<0.001, for comparisons between *LDLr^-/-^* and *WT* at specific days. For (B-G) Student’s T test, *P<0.05, **P<0.01, for comparisons between *LDLr^-/-^* and *WT*.

Though there are alternative mechanisms for colitis protection in the face of the dramatic increase in gut Tregs (such as other cell types or activities), we find no evidence of this in either *apoC-III^Tg^* or *LDLr^-/-^* mice. Given the significant increase in the number of Tregs that are accumulating in the *apoC-III^Tg^* intestine, and the fact that these are accompanied by a ∼100-fold increase in IL-10, we do not find it surprising that *apoC-III^Tg^* mice are protected from colitis despite no change to Treg activity/proliferation/suppressive capacity. Though multiple cell types secrete IL-10, these cells are not increased in *apoC-III^Tg^* mice during chow diet, Western diet, or after DSS-colitis induction (data not shown, no statistical differences between *WT* and *apoC-III^Tg^* lymphoid and myeloid populations in plasma, spleen, thymus, liver). Identifying the cell responsible for elevated IL-10 in *apoC-III^Tg^* mice is a critical issue that has yet to be resolved.

### Therapeutic treatment of apoC-III with chylomicrons protects from DSS-induced colitis

Our data strongly suggest that TAG transport processes are involved in Treg homeostasis and colitis in mice. We therefore hypothesized that the inhibition of TAG transport, but not apoC-III protein in the absence of lipoprotein TAG, would stimulate Tregs and colitis protection in *WT* mice. We used the DSS model of colitis in *WT* mice receiving a daily intraperitoneal injection of one of three treatments: 1) PBS vehicle control, 2) purified human apoC-III, or 3) purified human apoC-III bound to *WT* triglyceride-rich lipoproteins. ApoC-III was provided at a concentration five times higher than physiological levels (55 mg/dL) to mimic conditions of apoC-III overexpression [40]. Injections were given intraperitoneally to ensure the treatments are cleared through the intestinal lymphatics before entering into circulation [41]. Over the course of the treatment period, *WT* mice given apoC-III on lipoproteins (apoC-III + TRLs) lost significantly less body weight compared to the group that received apoC-III alone (\*\*\*\**P<0.0001*) (Figure 8A). When comparing the qualitative macroscopic damage associated with DSS treatment, *WT* mice given apoC-III + TRLs had a lower macroscopic damage score compared to apoC-III alone (Figure 8B). Finally, after the treatment period, gene expression analysis of the colon of DSS-treated mice revealed a distinct Treg signature in mice give apoC-III + TRLs (Figure 8C). These mice had upregulated expression of *FOXP3,* the master transcription factor for the Treg lineage and *IL-10*. These findings suggest that apoC-III on lipoproteins may provide protection from DSS-induced colitis through the accumulation of intestinal Tregs and their IL-10.

**Figure 8:**
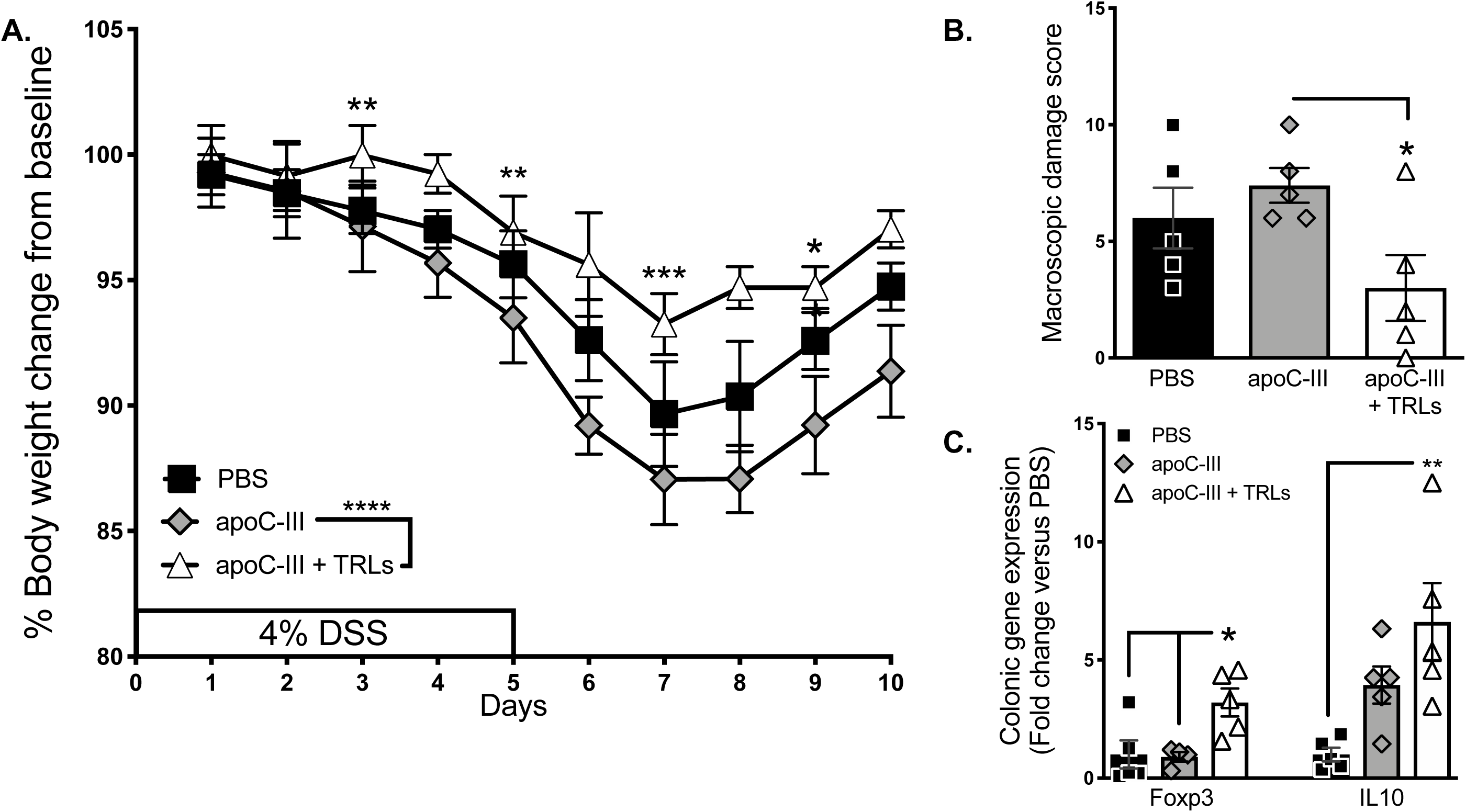
Therapeutic treatment of apoC-III with chylomicrons protects from DSS-induced colitis. *WT* mice (aged 8 weeks), both maintained on the same C57Bl/6J background and on chow diet, were given a daily injection of either PBS, purified apoC-III, or purified apoC-III + TRLs and treated with 4% DSS for 5 days. **(A)** Percent body weight changes from baseline (before treatment). **(B)** Macroscopic damage analysis of physiological symptoms of DSS. **(C)** Colonic mRNA expression after the complete treatment period (on Day 10). Results are expressed as mean ± SEM. n=5 per injection group. For (A), 2way ANOVA (****P<0.0001 for comparison between apoC-III and apoC-III+TRLs), with Sidak’s post-hoc analysis, *P<0.05, **P<0.01, ***P<0.001, for comparisons between injection treatments at specific days. For (B-C) Student’s T test, *P<0.05, **P<0.01, for comparisons between treatment groups.

## Discussion

Using genetic and *ex vivo* models that inhibit LDLr and triglyceride transport from lipoproteins into immune cells, we show that *apoC-III^Tg^* and *LDLr^-/-^* mice are both protected from DSS-colitis and simultaneously accumulate intestinal Tregs and IL-10. We show that T cells are capable of endocytosing TAG from lipoproteins, that this process is inhibited by apoC-III. This is the first evidence that Tregs and T cells use TAG, and that canonical regulators of TAG transport also regulate T cells and Tregs. We show that Tregs from *apoC-III^Tg^* are metabolically unique from *WT* Tregs and upregulate the genes of lipolysis and OXPHOS. Finally, we show that delivery of exogenous apoC-III on chylomicrons can protect *WT* mice from developing DSS-induced colitis.

A growing body of evidence suggests that Treg homeostasis relies on the ability to oxidize fatty acids. Tregs oxidize exogenous free fatty acids (palmitate), and the blockage of glycolysis, lack of exogenous glucose, or excess palmitate in the media will result in Treg enrichment, suppressive activity, and differentiation from CD4^+^ precursors[21,22,42,43]. In addition, *ex vivo* Tregs will take up exogenous FFAs at a greater rate when they are rapidly proliferating, and during activation [37,44]. Though these studies are critical to understanding how Tregs response to metabolic cues, they share the limitation that Tregs *in vivo* are normally exposed to triglyceride, not palmitate, in lipoproteins. We show for the first time that triglyceride-rich lipoproteins, the physiological lipid delivery system, can delivery Bodipy-TAG to T cells in culture, and that the loss of this pathway *in vivo* causes major changes to Treg accumulation, metabolism, and IL-10 concentrations in the gut. Physiologically, therefore, a key regulator of Treg function may be the presentation of triglyceride in lipoproteins.

Our data suggest that when Tregs are exposed to high levels of extracellular apoC-III on lipoproteins, those cells are deprived of exogenous lipid substrates. We find evidence for an intracellular switch in this condition, where *ex vivo* Tregs upregulate intracellular FAO but not glycolysis (via Seahorse and mRNA expression analyses). This finding is consistent with other studies that highlight Treg ability to switch from using TAG fuel to another other fuel sources which drive OXPHOS [38]. It will be important to determine how Tregs and T cells make up for the TAG they are unable to transport in the face of excess apoC-III. Prior to these findings, the role of extracellular triglyceride in immune cells was unknown.

Our study supports the findings by Rueda *et al.* that HDL internalization by Tregs promotes their survival and that blocking the oxidation of HDL lipids abolishes this effect [36]. Our studies add to this work by showing that lipoprotein triglyceride, and endocytosis, are critical for Tregs in models where LDLr is inhibited (both *apoC-III^Tg^* and *LDLr^-/-^* mice). Further, this mechanism of triglyceride metabolism/lipoprotein triglyceride endocytosis may be unique to intestinal immune cells; we only find Treg accumulation in the gut of *apoC-III^Tg^* and *LDLr^-/-^* mice. Further, by flow cytometry analyses of LDLr expression, Tregs can be stratified by their LDLr expression and their anatomical location, which Tregs in the gut expressing the highest LDLr. It is intriguing to speculate that the regulation by apoC-III might be unique to mLN Tregs, because these interact directly with intestinal lymph that carries diet-derived triglyceride and antigens on chylomicrons as well as intestinally-derived apoC-III. This would explain how apoC-III and LDLr can function as cardiovascular disease risk factors, where the thymic Treg responses predominate, and also act in an immunosuppressive manner in the gut [45].

Both *apoC-III^Tg^* and *LDLr^-/-^* mice are hyperlipidemic due to the inhibition of LDLr lipid uptake from plasma lipoproteins; therefore, we cannot fully dismiss a role for plasma hyperlipidemia in modulating tolerogenic gut T cells. However, the 12-week Western diet in *WT* mice which raised plasma triglyceride and cholesterol levels did not result in an accumulation of Tregs in the gut. This is consistent with several recent studies of plasma lipid metabolism and Treg biology. *ApoE^-/-^* mice produce dysfunctional splenic CD4^+^CD25^low^ Foxp3^+^ Tregs in response to a high fat diet [46]. In *LDLr^-/-^* mice, Tregs accumulate in the liver and migrate to the aorta during hypercholesterolemia but do not confer protection from atherogenesis [47]. A key difference in *apoC-III^Tg^* mice, however, is that unlike *apoE^-/-^* or *LDLr^-/-^* mice, which have very high plasma cholesterol levels, *apoC-III^Tg^* mice have significantly higher plasma triglycerides (∼1,500 mg/dL compared to ∼200 mg/dL in *LDLr^-/-^* and ∼150mg/dL in *apoE^-/-^)*. Hypercholesterolemia in *LDLr^-/-^* mice results in expanded Treg populations, but impedes their suppressive capacity [45]. It is possible that the different plasma lipids (cholesterol versus triglycerides) have varying roles in Treg homeostasis; our data would suggest that lack of access to this lipid also modifies Treg populations.

Our studies have not established the mechanism by which excess apoC-III causes Treg accumulation in the intestine. We find no differences between *WT* and Treg markers, nor are there changes to the basal IL-2 secretion in *apoC-III^Tg^* Tregs. Intestinal Treg accumulation in *apoC-III^Tg^* mice could also be due to an increase in Treg homing to the gut, an increase in T cell differentiation, a decrease in conversion of Tregs to anergic ex-Tregs, or another mechanism that results in a change in total numbers of Tregs. Overall, we find that apoC-III overexpression in Tregs does not influence their proliferation, suppressive capacity, anergy, or surface marker phenotype. *Ex vivo WT* and *apoC-III^Tg^* Tregs behave similarly, with no differences in any of these measured with TCR stimulation. This suggests that our *in vivo* finding of increased Tregs in the intestine is not due to a change in Treg activity/phenotype but is instead due to the accumulation of large numbers of Tregs in the gut. Our data characterizing Tregs supports this conclusion.

We can exclude the possibility that the *apoC-III^Tg^* mice are mounting an immune response against the *APOC3* transgene (thus resulting in the increase in intestinal Tregs) for multiple reasons. We do not observe any immune disease (intestinal or otherwise) in these mice, and they are immuno-competent since they do not succumb to any disease at a higher rate than their *WT* counterparts. They are able to mount a complete response to an intraperitoneal injection of lipopolysaccharide (data not shown). In addition, the highest expression of *APOC3* is in liver, and we find no change to liver-resident immune cell populations (Supplementary Figure 4), despite the high level of transgene expressed there. We also carried out an initial histological characterization of *apoC-III^Tg^* intestine and found no differences in the gross intestinal anatomy of these mice (small and large intestinal mucosa, infiltrating macrophages, intestinal length, size and number of intestinal villi) (data not shown). Our analysis of total plasma and spleen immune cells shows no underlying inflammation or depletion of parent cells, and there is no leakiness in intestinal permeability in the *apoC-III^Tg^* mice (assessed via FITC-dextran challenge). Furthermore, baseline analysis of whole blood and spleen shows no difference in T cell populations and monocytes, except for Tregs, suggesting that it is unlikely that these mice are mounting an immune response towards apoC-III.

Our findings highlight a novel role of lipoproteins and LDLr inhibition in stimulating intestinal tolerance and colitis protection in mice. Whether there are other critical lipoprotein receptors and apolipoprotein regulation (like glycosylphosphatidylinositol-anchored HDL binding protein 1 (GPIHBP-1), LDLr-related protein 1 (LRP1), or LPL) seems likely. Our data challenge the concept that lowering plasma triglycerides by inhibiting or by lowering apoC-III levels is universally beneficial; these data suggest that clinically, *APOC3* should be carefully considered in the context of both cardiovascular disease risk as well as IBD. Further, before treating patients with anti-apoC-III drugs, IBD risk may be an important clinical consideration. In short, for *APOC3*, what is good for the heart may not be good for the gut.

## Methods

### Animals

Male and female C57BL/6J (*WT*) and human apoC-III transgenic (*apoC-III^Tg^*) mice, both originating from heterozygous *apoC-III^Tg^* founders, were used for all studies (age 8-24 weeks). *ApoC-III^Tg^* mice were backcrossed six generations to C57Bl6/J mice. *ApoC-III^-/-^* mice (Stock #002057, originally generated by Maeda, et al. [48]) and *LDLr^-/-^* mice (Stock # 002207, originally generated by Hammer and Herz, et al.[49]) were purchased from The Jackson Laboratory (Bar Harbor, ME) and bred in-house. A separate set of *WT* controls were bred from heterozygous founders of the *apoC-III^-/-^* colony and *WT* mice (stock #000664) were purchased from The Jackson Laboratory to use as *LDLr^-/-^* controls. *ApoC-III^Tg^* mice (founders generously provided by Dr. H. Dong, University of Pittsburgh) were originally generated by Dr. Jan Breslow [26]. All baseline measurements were taken from mice maintained on chow diet with no treatment.

All strains of mice were housed in the same facility, 2-5 animals per cage, in a temperature-controlled (25 ±1°C) vivarium on a 10-h light-dark cycle. Unless noted, mice had free access to water and chow diet (LM-485 Mouse/Rat Sterilizable Diet, Harlan Laboratories). For Western diet experiments, *WT* mice received Western diet (TD.88137, Harlan, Envigo), containing 42% kcal of butterfat and 0.2% of cholesterol, for 12 weeks *ad libitum*.

### Blood measurements

After an overnight fast, blood via tail clip was collected from mice to isolate plasma. Plasma triglycerides and cholesterol were determined by chemical assay (Randox #TR210 and #CH200, respectively, UK), according to manufacturer’s protocol. For plasma cytokines, a terminal cardiac puncture was performed and whole blood was collected for serum isolation. Whole blood sat at room temperature for 20 minutes before centrifuging at 2,000 times gravity. Serum was isolated and frozen at −80°C until used for analysis. To determine IL-6, IL-10, IL-4, TNF-α, and IFN-γ levels, the EMD Millipore milliplex assay (catalog #MCYTOMAG-70k-05) was used according to the manufacturer’s protocol; for IL-10 and TNF-α, a BioLegend IL-10 ELISA assay (catalog #431411) and TNF-α ELISA assay (catalog #430901, San Diego CA) were used, respectively, according to the manufacturer’s protocol. For gut permeability analysis, mice were fasted for 6 hours and administered FITC-dextran (125 µg/µL, Sigma-Aldrich, St. Louis, MO) by oral gavage at 600 mg/kg body weight. After 1 hour, blood was collected for plasma analysis of FITC-dextran on a fluorescence spectrophotometer (BioTek Instruments, Inc., Winooski, VT) at excitation 485 nm and emission 535 nm. A standard curve was generated by diluting FITC-dextran in untreated mouse plasma.

### Induction of colitis

Two weeks prior to all experiments, dirty cage bedding was swapped between *apoC-III^Tg^*, *apoC-III^-/-^*, and *LDLr^-/-^* mice and their respective *WT* controls, to ensure identical environmental conditions[50]. Administration of 4-5% DSS (Colitis grade, M.W. 35 – 50 kDa, MP Biomedicals) was given *ad libitum* in drinking water for 5 days, followed by fresh water for 5 days post-DSS treatment. Mice were assessed daily for body weight changes and macroscopic damage score [50]. Colitis symptoms were scored (0 = normal, 1 = mild, to 3 = severe) for symptoms of hunched posture, ruffled fur, rate of breathing, crusty eyes, shivering, diarrhea, rectal bleeding, rectal prolapse, activity upon handling, and body weight change. Mice with a score greater than 3 were sacrificed, and all mice were sacrificed at day 10. To calculate macroscopic damage score, the sum of the total scores for the 10-day period was averaged for each genotype.

### ApoC-III treatment during colitis induction

*WT* Mice were randomly assigned a daily treatment of phosphate-buffered saline, purified apoC-III, and purified apoC-III with *WT* TRLs (n=5 per group). Treatments were delivered via intraperitoneal injection at the same time, daily for the ten-day DSS treatment period. To mimic conditions in *apoC-III^Tg^* mice, purified human apoC-III (Academy Bio-Medical Company) were given at a dose that is five times greater than healthy, physiologic levels of apoC-III (55 mg/dL per dose per mouse) [40]. ApoC-III with *WT* TRLs will be delivered at the same concentration with TRLs at 200 mg/dL to mimic the post-prandial state. Injection volumes were maintained at 200 μL.

### Tissue collection

Immediately following exsanguination using a terminal cardiac puncture, tissues were collected for flow cytometric or quantitative reverse transcription PCR (RT-qPCR) analysis. If collected for flow cytometry, tissue (MLNs, intestine, colon, spleen, and liver) was collected directly into phosphate buffered saline (PBS) and if collected for RT-qPCR (duodenum, ileum, colon, and liver) tissues were snap-frozen in liquid nitrogen and kept frozen at −80°C until RNA isolation. In addition, spleens were collected into PBS for Treg isolation.

### Preparation of single-cell suspensions for flow cytometric analysis

From the small intestine and the colon, the LP was isolated as previously described [51]. Briefly, small intestinal LP was isolated from the entire small intestine including duodenum through the ileum. The intestine and colon were incubated in predigestion buffer (HBSS with 5 mM EDTA and 1 mM DTT) for 30 minutes at 37°C, to remove epithelial cells, followed by two incubations in digestion buffer [collagenase Type 1 at 145 units/mL (Life Technologies) with DNase I at 25 units/µL (Invitrogen) in PBS], to break up connective tissue, releasing lymphoid cells. MLN, spleen, and liver were collected directly into PBS, and then tissue was mashed through a 70 µm strainer. MLN, spleen, and whole blood, which was collected via tail clip, were treated with erythrocyte lysis buffer (150 mM NH_4_Cl, 10 mM NaHCO_3_, 10 mM EDTA). All cells were pelleted and adjusted to 7-8 x10^6^ cells/mL.

### Antibody staining for flow cytometric analysis

Cells were stained with a live/dead discrimination dye (Zombie UV, catalog #423107, Biolegend). When appropriate, Fc receptors were blocked, using 10% FBS, before staining with anti-mouse fluorochrome-conjugated antibodies (Biolegend, see below for description). All antibody staining was conducted on ice, protected from light. Samples were fixed in 2% paraformaldehyde and permeabilized (1.6 mM Triton X-100 in PBS) for intracellular staining. For LDLr staining: cells were stained with primary LDLr antibody (Abcam, ab52818, 1:750), followed by secondary staining with Alexa 488 (Abcam, ab150065, 1:2,000). Cells were washed between each step using flow cytometry buffer (1x PBS, 5% FBS, 0.2 mM EDTA). Flow cytometry samples were run at the University of Connecticut Health Center Flow Cytometry Core. Samples were run using a BD LSR II (Becton-Dickinson Biosciences) after appropriate compensation. Collected events were set at 100,000.

Compensation controls using single stains were made using OneComp ebeads and UltraComp ebeads (Ebioscience) according to the manufacturer’s instructions. Automatic compensation was conducted by FACS Diva software. Fluorescence minus one (FMO) controls were used to determine appropriate gate positions and FMO samples were created using the same tissue as the experimental sample, at the same time of experimental preparation.

Our gating strategy is summarized as follows. When gating any sample, leukocytes were identified by forward scatter (FSC)-area (FSC-A) by side scatter (SSC)-area, followed by identification of single cells using FSC-height by FSC-width. Finally, live cells were gated on using FSC-A by Live/Dead viability dye. Absence of the dye indicates live cells. Live cells were then identified as T cells using CD3 by FSC-A, and then further identified as helper T cells with CD4 by FSC-A. To distinguish Tregs, CD25 was gated by FOXP3 and double-positive cells were selected (CD25^+^FOXP3^+^). FMO controls were used to determine all gates except FSC by SSC and single cells.

FlowJo software (Treestar, Ashland, OR) was utilized for final analysis of collected data. For absolute cell numbers, the percentage of living cells of a certain subset was multiplied by the number of living cells as determined by a TC20 Automated Cell Counter (Bio-Rad Laboratories, Inc. Hercules, CA).

The following Biolegend antibodies were used: Brilliant Violet 711 CD3ε (Clone: 145-2C11), PerCP-Cy5.5 CD4 (Clone: RM4-5), APC-Cy7 CD8a (Clone: 53-6.7), Alexa Fluor 700 CD19 (Clone: 6D5), Brilliant Violet 510 CD14 (Clone: Sa14-2), PE-Cy5 CD25 (Clone: PC61), PE FOXP3 (Clone: MF-14), APC CD11c (Clone: N418), Alexa Fluor 700 I-A/I-E (Clone: M5/114.152), Brilliant Violet 421 CD103 (Clone: 2E7), Alexa Fluor 594 Ki67 (Clone: 11F6), and APC-Cy7 IL-10 (Clone: JES5-16Eep). APC anti-CCR9 (Clone: CW-1.2) was purchased from Life Technologies. Cells are putatively defined as following: CD3^+^CD4^+^CD25^+^FOXP3^+^ for Tregs, CD11c^+^MHC^+^CD103^+^ for tolerogenic DCs and CD3^-^CD14^+^ for monocytes.

### Tag-It Violet™ proliferation analysis

For proliferation analysis, cells were stained with Tag-It Violet™ Proliferation and Cell Tracking Dye according to the manufacturer’s instructions (Catalog #425101, Biolegend, San Diego, CA). Briefly, cells were stained for 12 minutes at 37°C with a 5 µM solution of Tag-It Violet™ in RPMI-1649 media without serum. Staining was quenched with five times the volume of RPMI-1640 with 10% FBS. Cells were then utilized for downstream applications. Proliferation analysis was conducted using FlowJo’s Proliferation Tool (Treestar, Ashland, OR). Proliferation index is defined as the total number of divisions divided by the number of cells that went into division.

### T-cell isolation and culture

Tregs, defined CD4^+^CD25^+^, were isolated from the spleen and lymph nodes of *WT*, *apoC-III^Tg^*, and *apoC-III^-/-^* mice using positive selection with MojoSort Streptavidin Beads (Catalog #480016, Biolegend) and Biotin-conjugated antibodies according to the manufacturer’s instructions (anti-CD4, clone: GK1.5 and anti-CD25, clone: PC61). Briefly, single-cell suspensions from spleen were generated and cell concentrations were adjusted to 10^6^ cells/mL in PBS. Samples were incubated for 20 minutes with 10 µL of CD4^+^ antibody followed by incubation with streptavidin-conjugated beads. MojoSort Magnet (BioLegend, catalog #480019) was used for two collections of the positive fraction and the collected CD4^+^ cells were used for the same staining and collection process with anti-CD25 antibody (for Tregs). To determine cell purity, an aliquot of cells was stained with an anti-CD4 (clone: Rm4-4) and anti-Foxp3 antibody. Cells had 80% purity or greater when gated on CD4 and Foxp3. Cell purity was determined using the gating strategy shown in Supplementary Figure 6A and FMOs for CD4 and Foxp3 are shown in Supplementary Figure 6B. Isolated cells were used for RNA isolation by freezing in Trizol or were plated in complete RPMI-1640 media for proliferation analysis or Seahorse Mito Stress Analysis. When cultured, purified IL-2 (25 U/mL), anti-CD3 (0.5 µg/mL), and anti-CD28 (1 µg/mL) (Biolegend, catalog #504701, 100201, and 102101) were added to maintain a CD4^+^CD25^+^ Treg population. *WT* CD4^+^CD25^+^ Tregs cultured for proliferation analysis were treated with TRLs from apoC-III^-/-^, *WT,* and apoC-III^Tg^ mice for 96 hours. Plasma was standardized by triglyceride content after a triglyceride assay.

### *Ex vivo* Treg analysis

CD4^+^CD25^+^ cells were isolated using negative selection of CD4^+^ T cells using the MojoSort Mouse CD4 Naïve T-cell Isolation Kit (catalog #480040, Biolegend) followed by positive selection of CD25^+^, as described above. Isolated cells were cultured in complete media with and without TCR stimulation. For TCR stimulation, anti-CD3 (0.5 µg/mL) and anti-CD28 (1 µg/mL) were added to complete RPMI media. After 24 hours, cells were collected and stained with following phenotyping markers: Brilliant Violet CD152/CTLA-4 (Clone: UC10-4B9), PE CD223/LAG3 (Clone: C9B7W), PE/Cy7 CD357/GITR (Clone: YGITR765), PerCP/Cy5.5 CD279/PD-1 (Clone: 29F.1A12), APC CD304/Neuropilin-1 (Clone: 3E12), and Alexa Fluor 594 Ki67 (Clone: 11F6). Cell culture supernatant was collected and frozen for IL-2 analysis using LEGENDplex*™* technology according to the manufacturer’s instructions (Biolegend).

### Isolation of triglyceride-rich lipoproteins

After a four-hour fast, *WT* and *apoC-III^-/-^* mice were gavaged with Bodipy FL C12 (D3822, Life Technologies, Eugene, OR) at 2 µg/g body weight dissolved in olive oil or olive oil alone at a final volume of 200 µL. Three-four hours post-gavage, blood was collected via cardiac puncture and allowed to coagulate at room temperature. Serum was isolated via centrifugation at 2,000 x gravity for 20 minutes. Serum was transferred to a 4.7 mL ultracentrifuge tube and overlaid with 300 µL saline. Samples were spun at 50,000 rpm in a TLA110 rotor using the Optima Max-TL Ultracentrifuge (Beckman Coulter) for over 12 hours, and the top fraction was collected for use. Samples containing Bodipy were protected from light throughout preparation. Isolated TRLs were used in Jurkat T-cell experiments, cell proliferation analyses, and Seahorse analyses.

### Jurkat T-cell cultures

Jurkat T-cells (clone E6-1, ATCC TIB-152, Manassas, VA) were grown in RPMI-1640, supplemented with 10% FBS and 1% penicillin/streptomycin in an atmosphere of 5% CO_2_ at 37°C. When treated with TRLs, cells were serum-starved overnight then administered TRLs with and without purified human apoC-III at 20 µg/mL (Academy Bio-Medical Co, Houston, Texas). For lipid uptake experiments, cells were treated for 48 hours, and then collected for flow cytometric analysis. For proliferation analysis, cells were stained with Tag-It Violet Proliferation dye (catalog #425101, Biolegend), treated for 96 hours, and then collected for flow cytometric analysis. For RNA analysis, cells were collected in Trizol.

### Seahorse XF Cell Mito stress analysis

The OCR was measured with an XFp analyzer (Agilent Technologies, Santa Clara, CA). Cultured primary CD4^+^CD25^+^ Tregs were isolated and cultured as described above. Isolated cells were incubated for 12-16 hours in RPMI-1640 media, and then cells were seeded at a density of >200,000 cells per well of an XFp cell culture microplate. Before assay, cells were equilibrated for 1 h in unbuffered XF assay medium supplemented with 1 mM pyruvate, 2mM glutamine and 10 mM glucose. Mixing, waiting, and measure times were 2, 2, and 4 min, respectively. Compounds were injected during the assay at the following final concentrations: 7.5 µM oligomycin and 7.5 µM FCCP, for OCR measurements.

### qRT-PCR

Snap-frozen organ fragments were mechanically homogenized with the use of a Tissue Tearor (Biospec Products, LLC). Total RNA was extracted from homogenized tissue and isolated cells using Tri-Reagent (LifeTechnologies) according to the manufacturer’s instructions. cDNA was synthesized using a Reverse Transcriptase kit (Promega). Quantitative real-time PCR was conducted using primer sets in Supplementary Table 1 and mRNA levels were analyzed in duplicate using SYBR green (Invitrogen) using a thermocycler (BioRad CFX Connect Real Time System). Samples were normalized to cyclophilin. For fold change, samples were then standardized to *WT* averages.

### Statistics

Statistical analyses were performed using GraphPad Prism software, version 6. All results are expressed as mean ± SEM. Results were analyzed by Student’s *t* test (unpaired with equal standard deviations), comparing genotypes or treatments, after a test to identify outliers was conducted (ROUT outliers, Q=1%) or using a two-way ANOVA with Sidak’s post-hoc analysis after outliers were identified. Differences with a *P* value of less than 0.05 were considered statistically significant (\**P<0.05, **P<0.01, ***P<0.0001, ****P<0.00001)*.

## Data Availability

The data that support the findings in this study are available from the corresponding author upon request.

## Study Approval

All animal procedures were performed in accordance with the University of Connecticut Internal Animal Care and Use Committee and in compliance with the National Institutes of Health Guide for the Care and Use of Laboratory Animals.

## AUTHOR CONTRIBUTIONS

C.N.R. and A.B.K designed experiments and wrote and prepared the manuscript. C.N.R., D.L., Z.K.J., and N.S.T performed experiments. E.R.J. designed and oversaw all flow cytometry experiments. C.N.R. and A.B.K analyzed the data and constructed the figures. E.R.J., A.T.V., and A.B.K provided insight and edits for manuscript preparation.

## ACKNOWLEDGEMENTS

This work was funded by grants to ABK from NIH (DK101663, DK116011, DK118239).

## CONFLICTS OF INTEREST

The authors have no conflicts of interest to report.

---

**Supplementary Figure 1:**
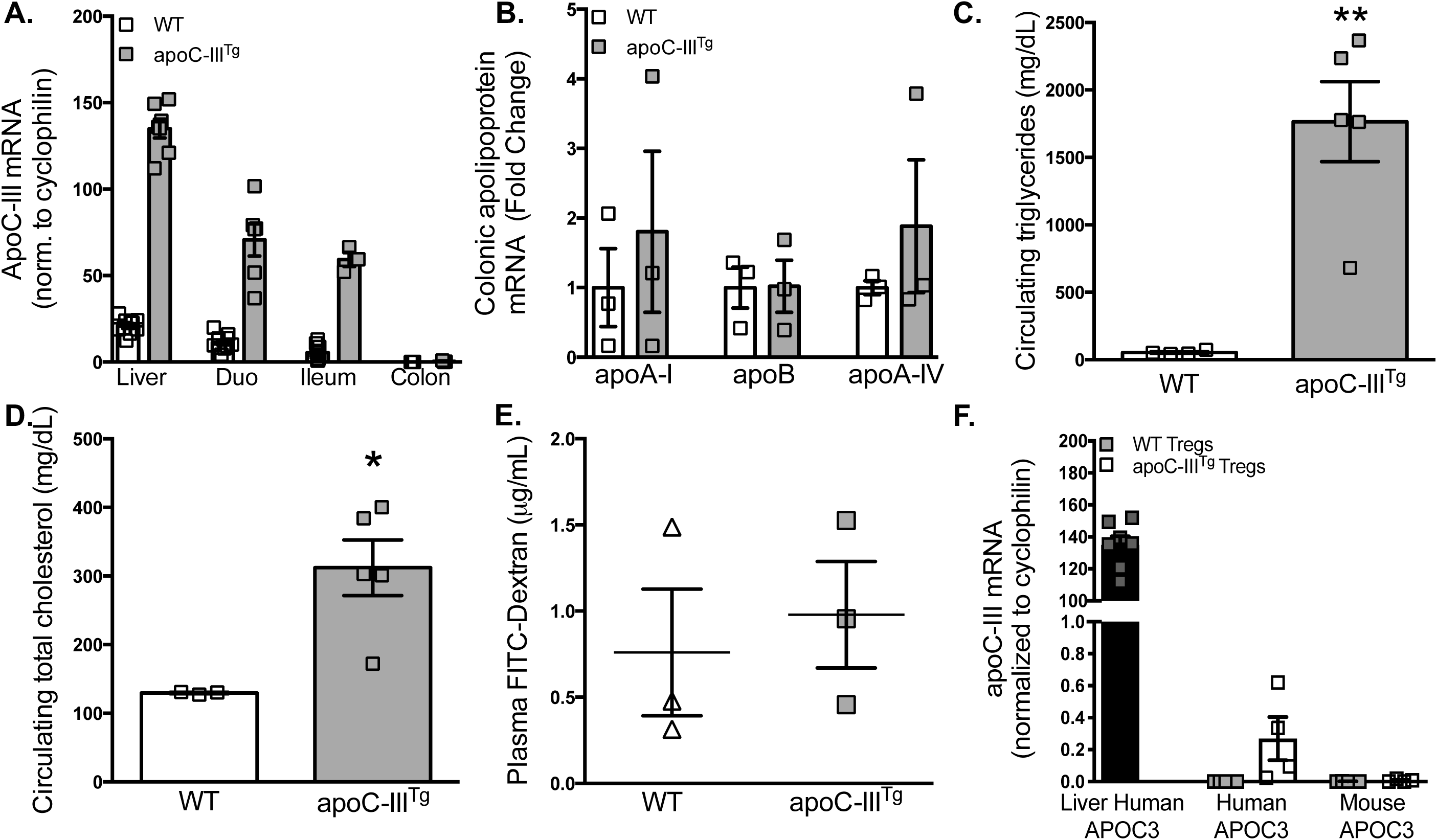
*ApoC-III^Tg^* mice have a baseline hyperlipidemic phenotype that results from overexpression of human APOC3. **(A)** mRNA expression of APOC3 in the liver, intestine and colon of *WT (*endogenous APOC3) and *apoC-III^Tg^* mice (human APOC3 transgene). (n=3-10) **(B)** Colonic expression of apolipoproteins (apoA-I, apoB and apoA-IV) in *WT* and *apoC-III^Tg^* mice. (n=3) **(C-D)** Fasted plasma triglycerides (C) and cholesterol (D) of *apoC-III^Tg^* mice compared to *WT*. (n=3-8) **(E)** Plasma levels of FITC-labeled Dextran, post-oral gavage, of *WT* and *apoC-III^Tg^* mice. **(F)** APOC3 (human transgene and mouse) mRNA expression from Liver *WT* and *apoC-III^Tg^* Tregs normalized to cyclophilin. Results are expressed as mean ± SEM. Student’s T test, *P<0.05, **P<0.01

**Supplementary Figure 2:**
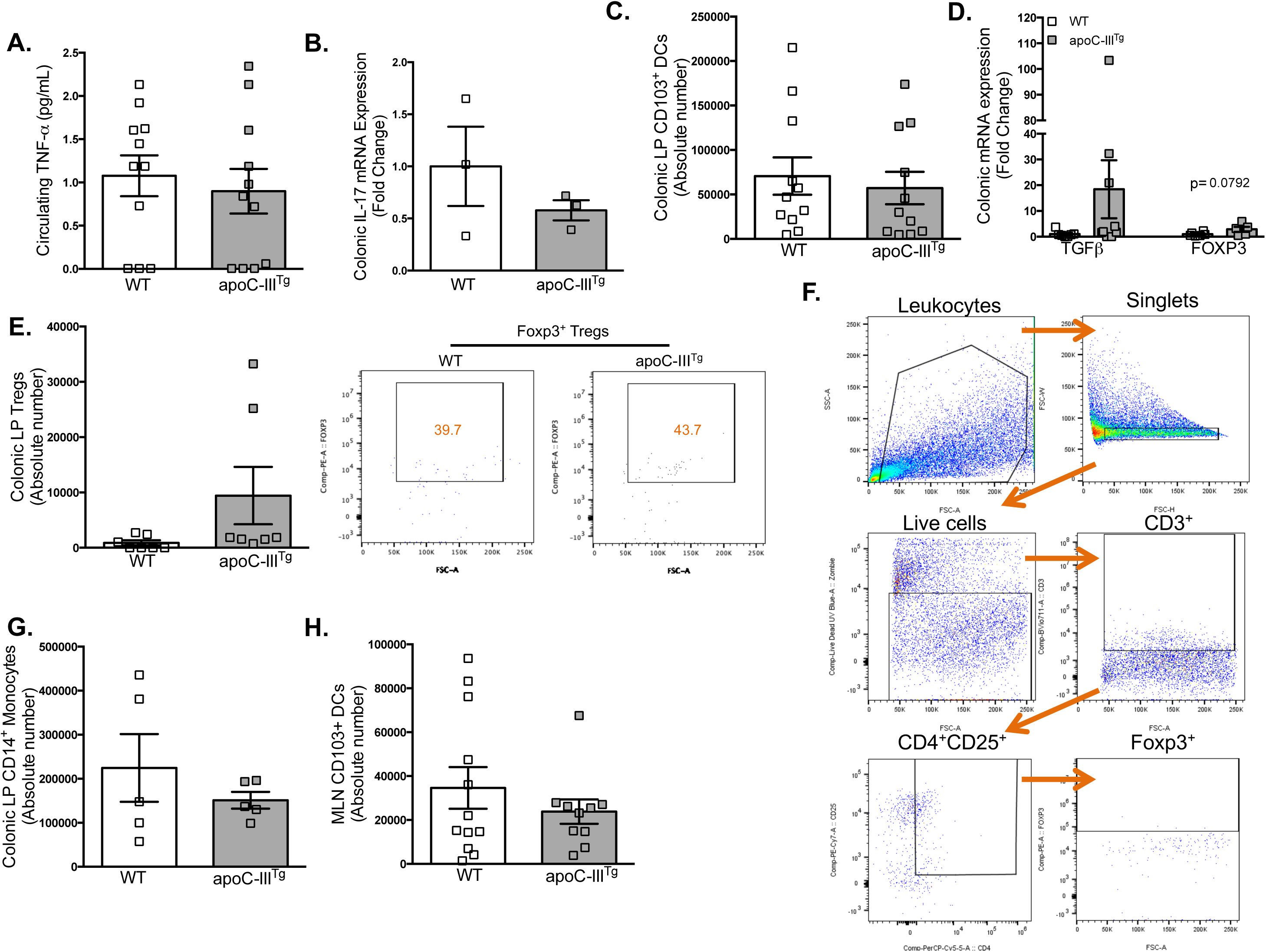
ApoC-III^Tg^ mice maintain gut tolerance after DSS treatment. *ApoC-III^Tg^* and *WT* mice (aged 8 weeks), both maintained on the same C57Bl/6J background and on chow diet, were treated with 4% DSS for 5 days followed by 5 days of water. After the 10 day treatment period, we analyzed **(A)** circulating TNF-α concentration, **(B)** colonic IL-17 mRNA expression, **(C)** flow cytometric analysis of the absolute number of colonic LP CD103^+^ DCs. (n=11) **(D)** Colonic mRNA Expression of Treg-related genes, *TGF-β* and *FOXP3*. (n=5-6) **(E)** Colonic LP Tregs. (n=7) **(F)** Flow cytometry gating strategy to identify colonic LP Tregs. **(G)** Colonic LP CD14^+^ monocyte. (n=5) and **(H)** MLN CD103^+^ DCs. (n=10-12). Results are expressed as mean ± SEM, from three independent DSS treatments.

**Supplementary Figure 3:**
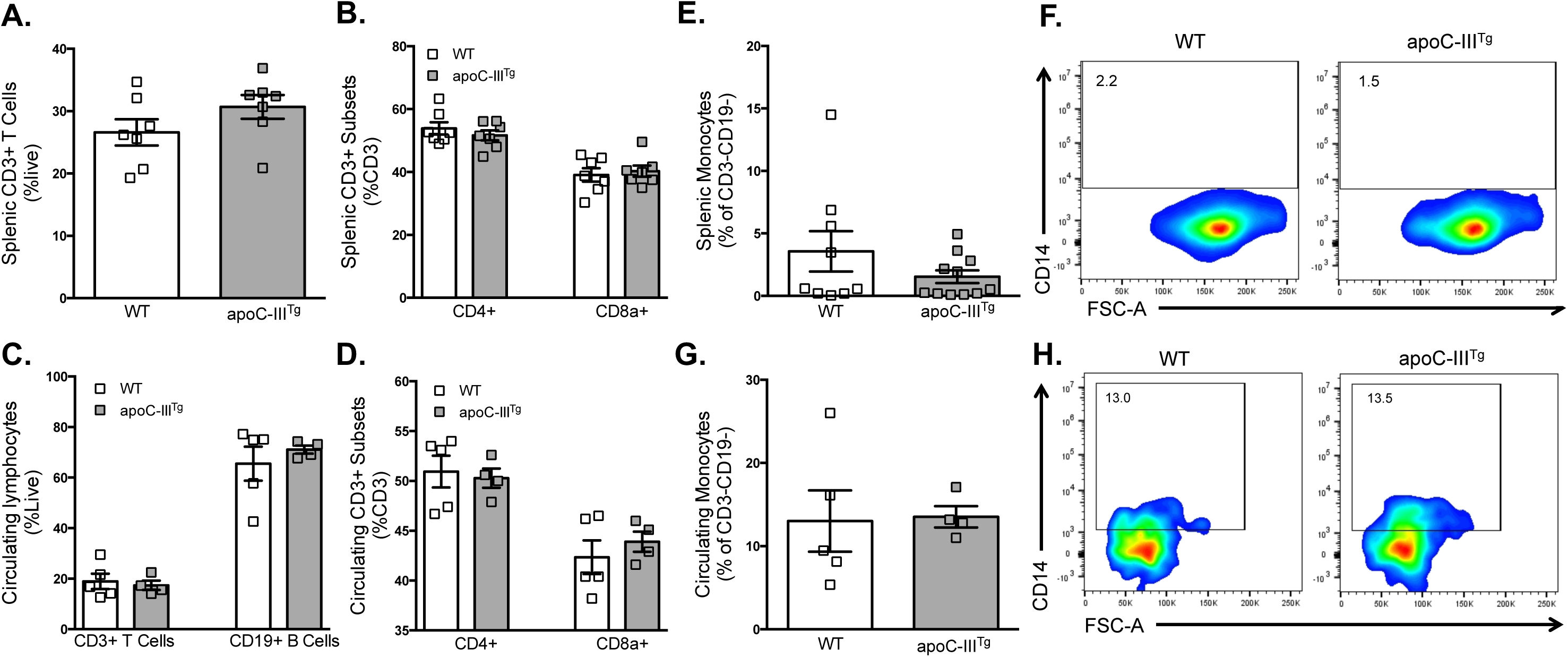
ApoC-III^Tg^ mice have no changes in splenic and circulating immune subsets. **(A-D)** Splenic (A-B) and circulating (C-D) CD3^+^ T cells, CD3^+^ subsets, and CD19^+^ B cells. **(E-H)** Splenic (E-F) and circulating (G-H) CD14^+^ monocytes in *WT* or *apoC-III^Tg^* mice. Dot plots are representative of three experiments. Results are expressed as mean ± SEM.

**Supplementary Figure 4:**
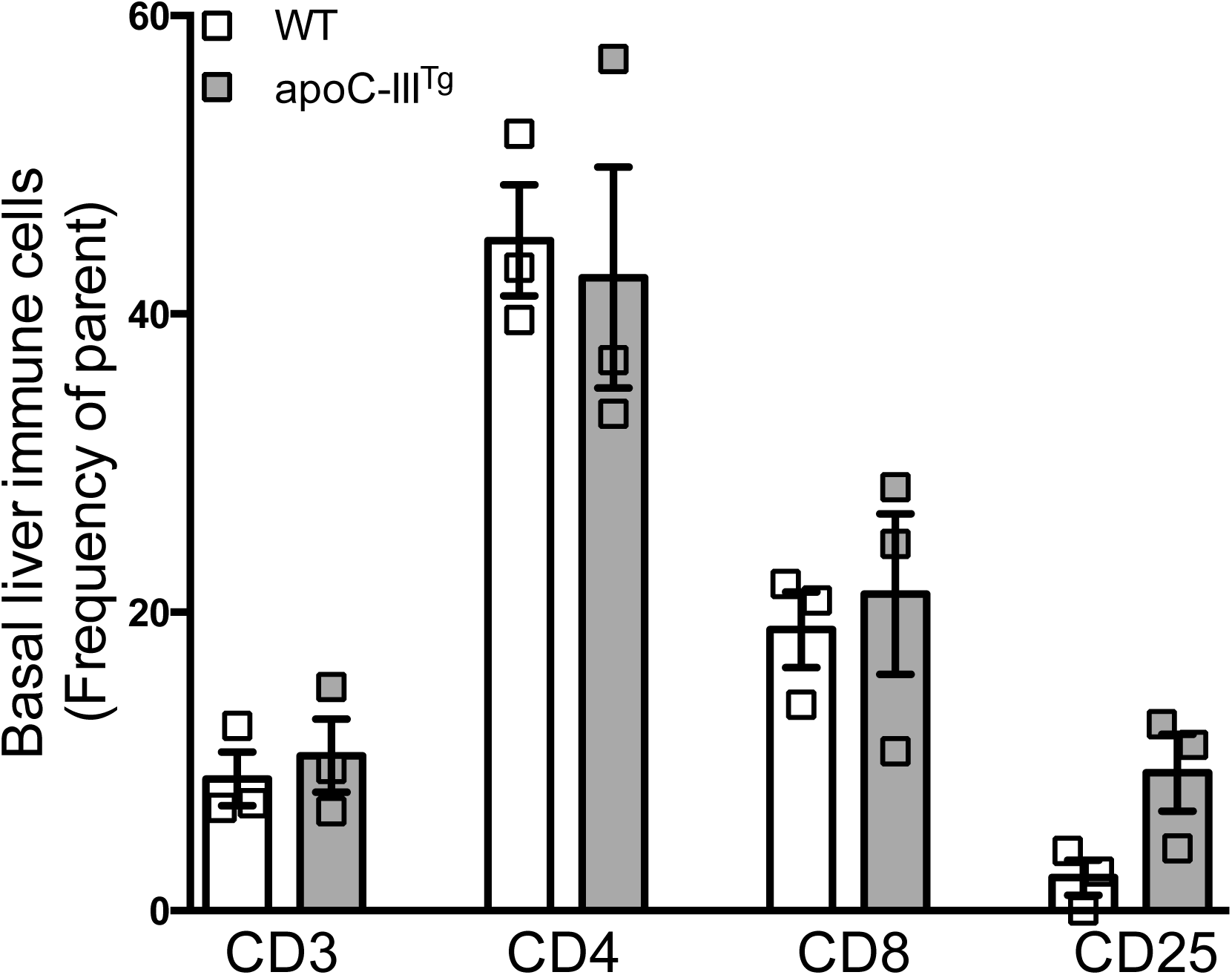
Hepatic baseline immune cell profile of WT versus apoC-III^Tg^ mice, analyzed by flow cytometry.

**Supplementary Figure 5:**
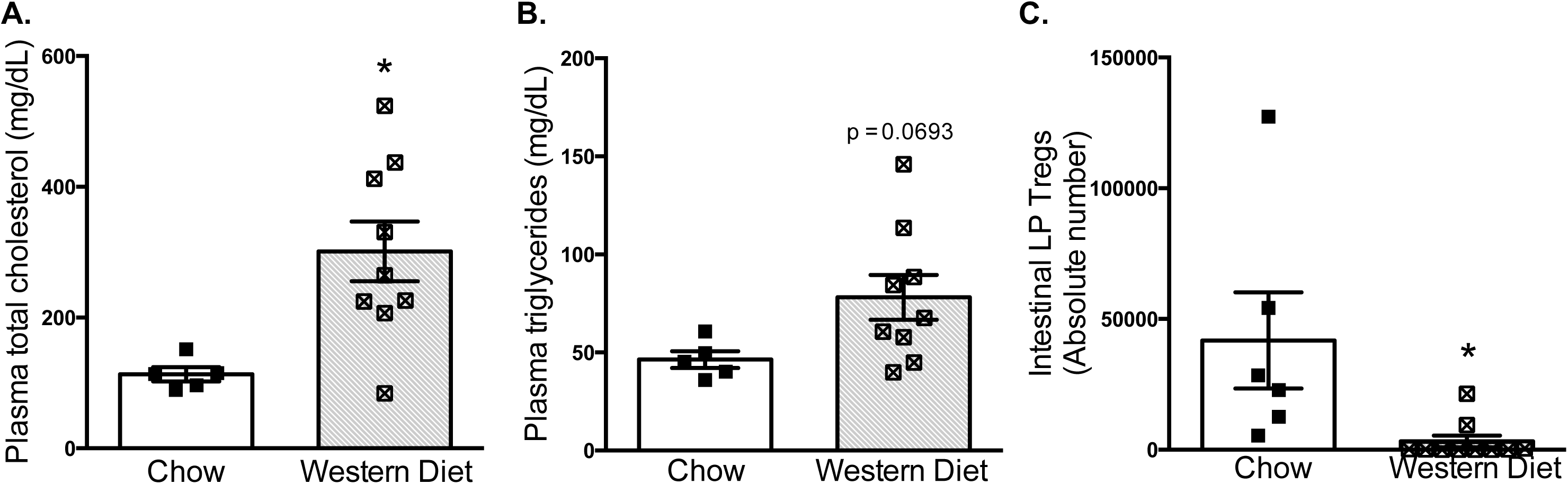
Hyperlipidemia does not promote accumulation of intestinal Tregs in WT mice. **(A-B)** Fasting plasma total cholesterol (A) and triglycerides (B) after 4 weeks of a high-fat, high-cholesterol Western diet. **(C)** Flow cytometric analysis of Tregs from intestinal LP from *WT* mice fed a Western diet for 12 weeks. N=5-9. Results are expressed as mean ± SEM. Student’s T test, *P<0.05.

**Supplementary Figure 6:**
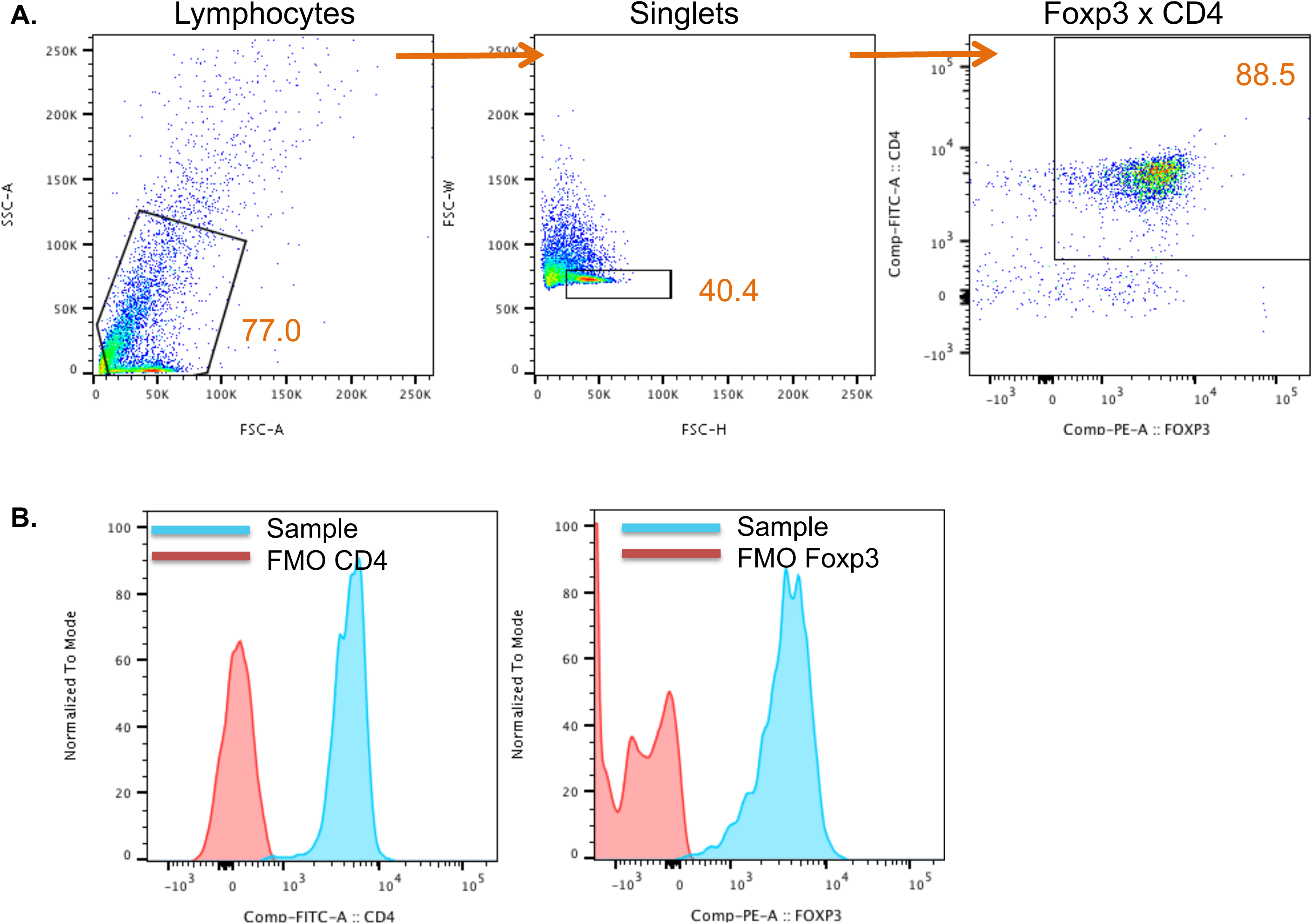
Sample purity check for isolated Tregs using flow cytometric analysis. Isolated CD4^+^CD25^+^ cells were gated using FSC-A by SSC-A to determine size followed by FSC-H by FSC-W to identify single cells. Final Treg gate was back-gated to ensure CD4^+^Foxp3^+^ cells were definitively populations of interest and not debris. **(A)** To determine purity of isolated cells, Foxp3 was gated by CD4. Tregs populations were greater than 80% pure. **(B)** FMOs for CD4 and Foxp3 were used to determine CD4^+^Foxp3^+^ gate. Sample histograms with FMO controls are shown for each marker.

**Supplementary Figure 7:**
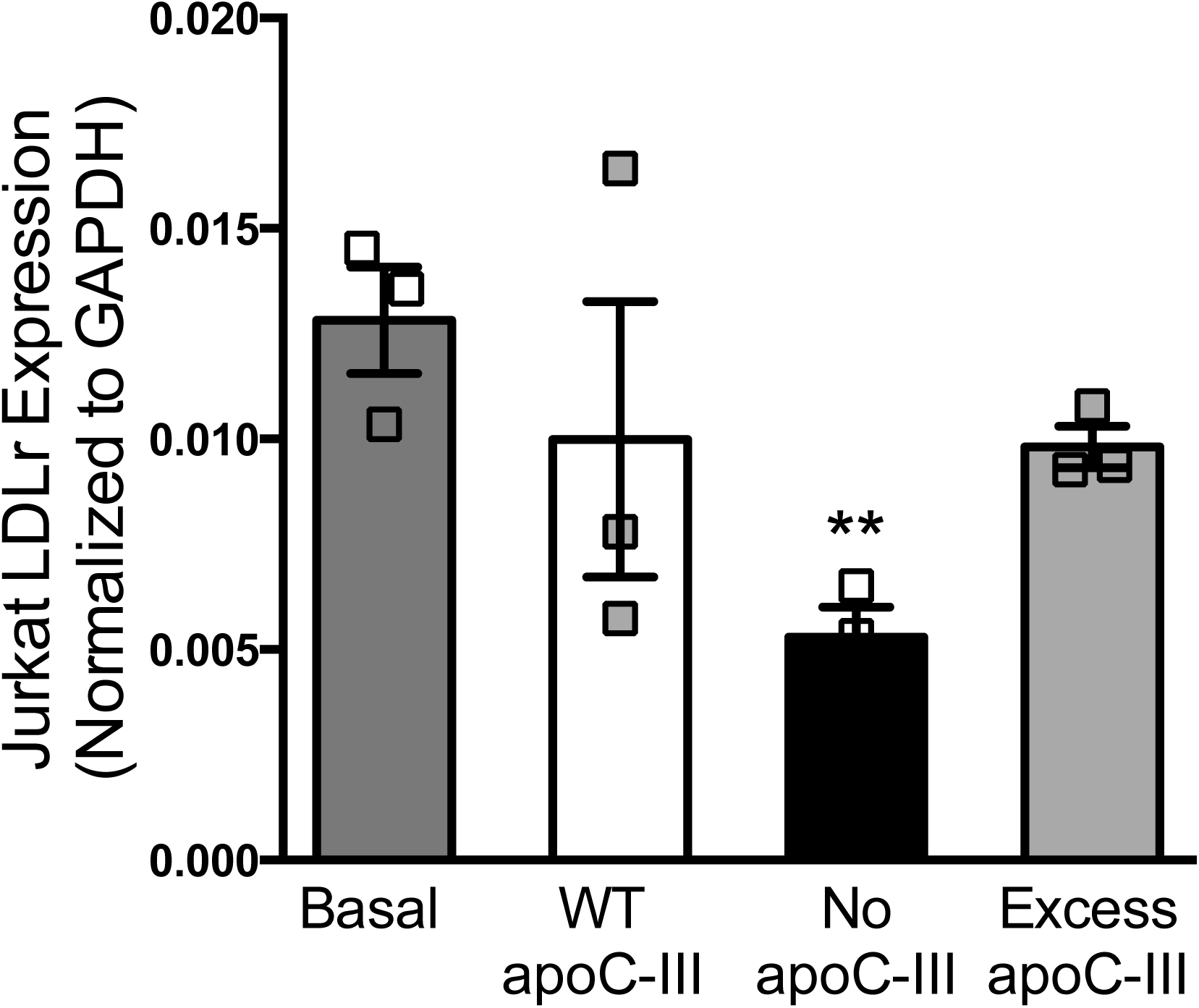
Jurkat T cells treated with apoC-III have comparable expression levels of LDLr to untreated cells. Jurkat T cell expression of LDLr in response to treatment with saline (Basal), WT TRLs (normal apoC-III), TRLs with human apoC-III (Excess apoC-III) and apoC-III^-/-^ TRLs (no apoC-III). Results are expressed as mean ± SEM. Student’s T test, **P<0.01 versus Basal control.

**Supplementary Figure 8:**
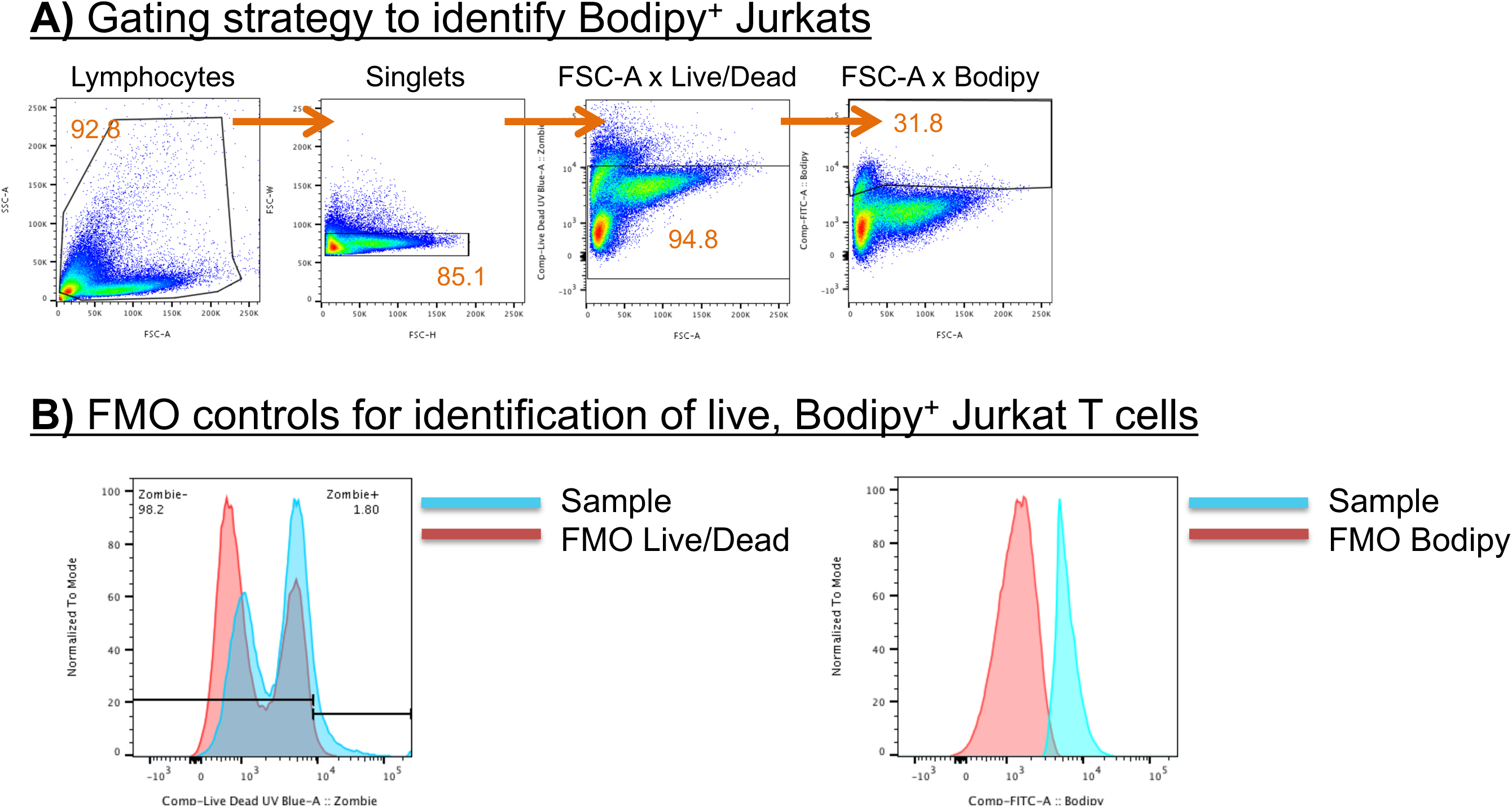
Sample gating strategy for identification of Bodipy^+^ Jurkat T cells. **(A)** Jurkat T cells were first identified by a lymphocyte gate from FSC-A x SSC-A followed by identification of singlets from FSC-H x FSC-W. Live cells were identified using an FMO for Zombie Live/Dead Discriminate Dye. Since Live/Dead Discriminate Dye stains dead cells, all cells shown in FMO Live/Dead (without dye) are live and selected as our gate of interest. **(B)** Bodipy^+^ cells were identified using FMO Bodipy.

**Supplementary Figure 9:**
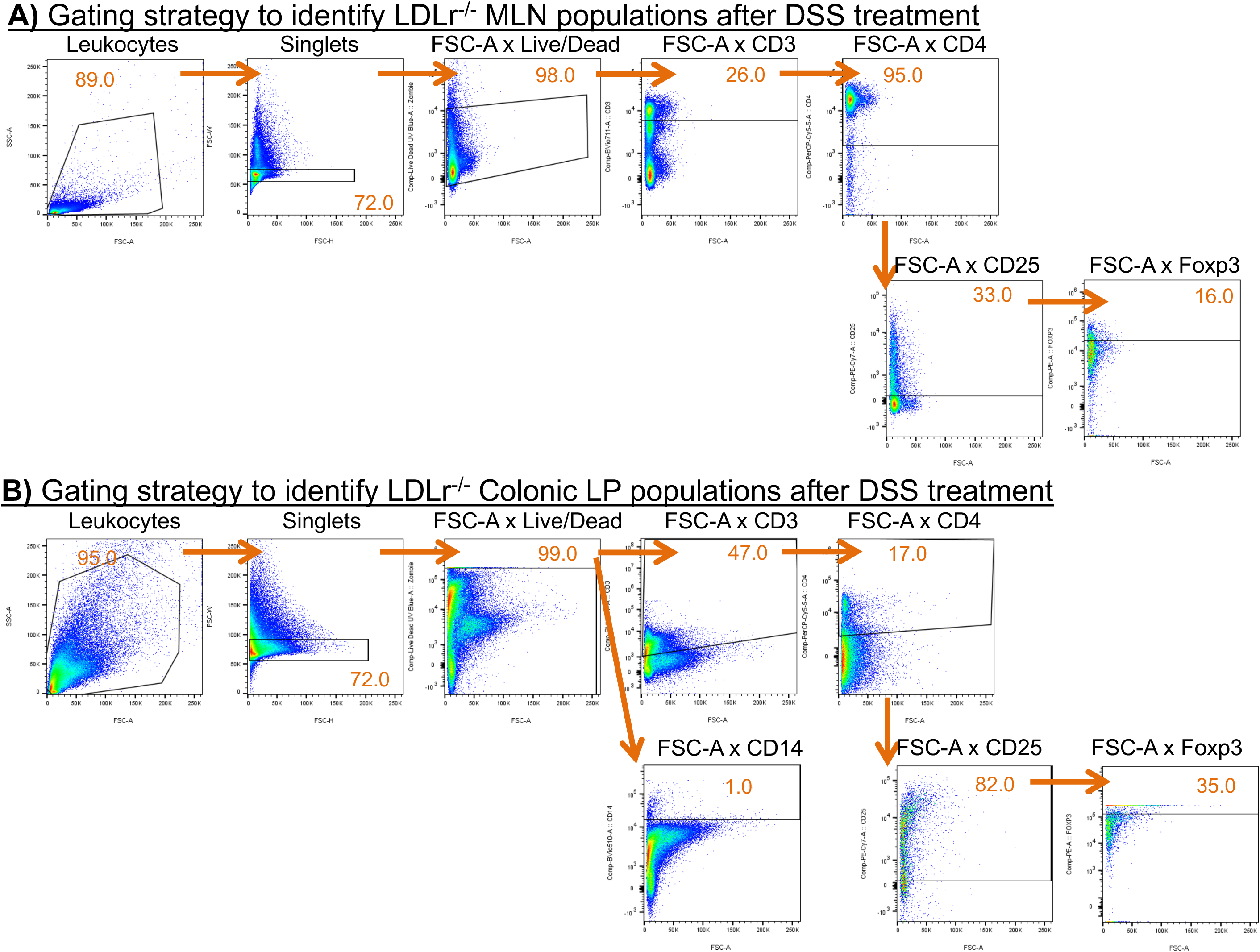
Sample gating strategy for flow cytometric analysis from data in Figure 7. **(A)** Complete gating strategy for identification of MLN CD4^+^ T cells (top row) and Tregs (bottom row) after DSS treatment. **(B)** Complete gating strategy for identification of colonic LP CD4^+^ T cells (top row) and Tregs (bottom row) after DSS treatment.

**Supplementary Table 1:**
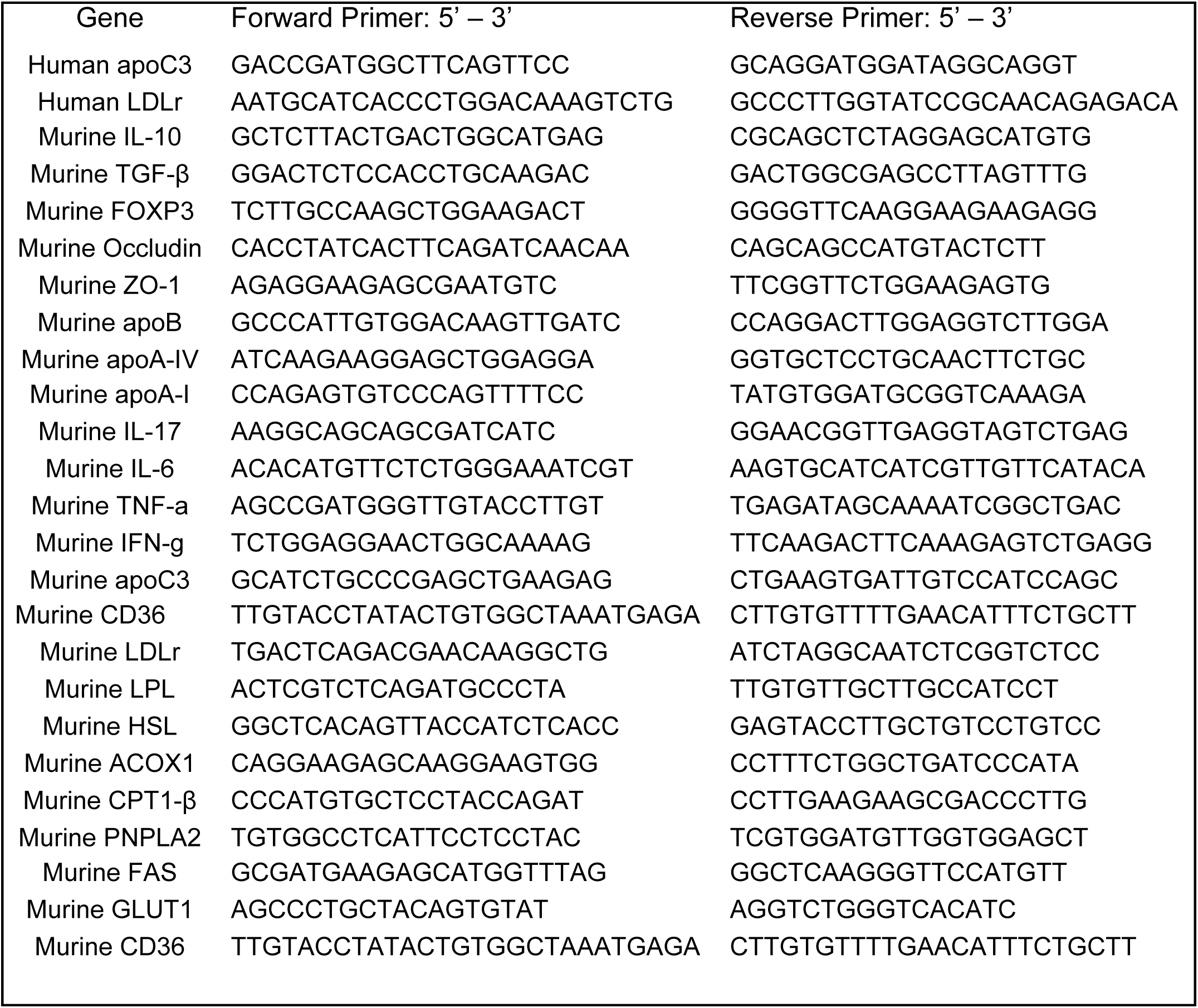
mRNA primers for RT-qPCR analysis.

